# The dimerized pentraxin-like domain of the adhesion G protein-coupled receptor 112 (ADGRG4) suggests function in sensing mechanical forces

**DOI:** 10.1101/2022.09.20.508530

**Authors:** Björn Kieslich, Jana Brendler, Albert Ricken, Torsten Schöneberg, Norbert Sträter

**Author notes:** Correspondences should be addressed to: Nobert Sträter, Institute of Bioanalytical Chemistry, Center for Biotechnology and Biomedicine, Leipzig University, 04103 Leipzig, Germany, Torsten Schöneberg, Rudolf Schönheimer Institute of Biochemistry, Medical Faculty, Leipzig University, Johannisallee 30, 04103 Leipzig, Germany.

## Abstract

Adhesion G protein-coupled receptors (aGPCRs) feature large extracellular regions (ECRs) with modular domains that often resemble protein classes of various function. The pentraxin (PTX) domain, which is predicted by sequence homology within the ECR of four different aGPCR members, is well known to form pentamers and other oligomers. Oligomerization of GPCRs is frequently reported and mainly driven by interactions of the seven-transmembrane region and N- or C-termini. While the functional importance of dimers is well-established for some class C GPCRs, relatively little is known about aGPCR multimerization. Here, we showcase the example of ADGRG4, an orphan aGPCR that possesses a PTX-like domain at its very N-terminal tip, followed by an extremely long stalk containing serine-threonine repeats. Using x-ray crystallography and biophysical methods we determined the structure of this unusual PTX-like domain and provide experimental evidence for a homodimer equilibrium of this domain which is Ca^2+^-independent and driven by intermolecular contacts that differ vastly from the known soluble PTXs. The formation of this dimer seems to be conserved in mammalian ADGRG4 indicating functional relevance. Our data alongside of theoretical considerations lead to the hypothesis that ADGRG4 acts as an *in vivo* sensor for shear forces in enterochromaffin and Paneth cells of the small intestine.

## Introduction

Although it is generally accepted that G protein-coupled receptors (GPCRs) exist as monomers there is convincing evidence that some GPCRs can also form dimers and even higher-order oligomers. Members of the metabotropic glutamate receptor-like GPCRs (class C), are well-known examples for forming stable homo- and heterodimers (Møller et al. 2018). Moreover, transient oligomerization was frequently reported for rhodopsin-like (class A) receptors (Walsh et al. 2018; Milligan et al. 2019). In class B, the secretin receptor was one of the first members for which dimerization was described (Ding et al. 2002), yet other reports about homo- and heterooligomerization followed (Ng and Chow 2015). Within the class B, adhesion GPCRs (aGPCRs) form a structurally separate receptor cluster comprising 33 members in the human genome which can be further subdivided into nine families (Hamann et al. 2015). Here, homo- and heterodimerization was shown in *trans* and *cis*, mediated via their large extracellular regions (ECRs). For example, members ADGRC1–3 undergo homophilic *trans* interactions (Shima et al. 2007; Nishimura et al. 2012). ADGRE5 heterodimerizes with lysophosphatidic acid receptor 1 in prostate and thyroid cancer cell lines and thus amplifies RhoA activation (Ward et al. 2011; Ward et al. 2013).

There is a backlog demand for fundamental research on the aGPCR class, as the majority of members are still considered orphan receptors. However, there is an evident involvement of aGPCRs in various developmental and physiological processes and inherited diseases (Schöneberg and Liebscher 2021). Members of the aGPCR class often stand out with extraordinary long ECRs, which comprise module-like domain arrangements and many glycans. The so-called pentraxin (PTX)-like domain is one of these extracellular domains (ECDs). Sequence homology predicts PTX-like domains to be present within the ECRs of the aGPCR members ADGRD1 (GPR133), ADGRD2 (GPR144), ADGRG4 (GPR112), and ADGRG6 (GPR126) (Hamann et al. 2015). However, knowledge is lacking about the function of this domain within the ECR of aGPCRs. The term “pentraxin” is inevitably linked to the homopentamerization of the archetypal serum proteins C-reactive protein (CRP) and serum amyloid component P (SAP). Indeed, pentamerization and other oligomeric states beyond dimers have been reported for other pentraxins (Kirkpatrick et al. 2000; Inforzato et al. 2008). Therefore, the question arises whether PTX-like domains in ECRs of aGPCRs oligomerize in a similar fashion as their soluble counterparts. Furthermore, CRP and SAP feature distinct Ca^2+^-binding sites that are mandatory for ligand binding and, in case of CRP, for pentamerization (Pepys et al. 1977; Pepys 2018). Despite being distinct protein classes, the basic β-sandwich fold of the pentraxins is also shared by other domain classes. Specifically, the laminin G-like (LG) domains exhibit a high structural resemblance to PTXs, despite a low sequence similarity. A more distant structural homology is also found for legume lectins, galectins and bacterial β-glucanases (Rudenko et al. 2001). The laminin α subunit possesses five LG domains at its C-terminus (LG1–5), which contain the major adhesive sites of laminins in the basal membrane (Hohenester and Yurchenco 2013; Hohenester 2019). After discovery in the laminin α subunit (Hohenester et al. 1999), LG domains were identified in other proteins of diverse biological function (Rudenko et al. 2001). Their function was linked to e.g. α-dystroglycan binding and similar binding modes were suggested for the LG3 domains of the heparan sulfate proteoglycans agrin (Stetefeld et al. 2004) and perlecan (Le et al. 2011). In neurexins, LG domains mediate a Ca^2+^-dependent interaction with neuroligin (Chen et al. 2008), forming a synaptic adhesion complex. LG domains were identified in the sex-hormone binding globulin (SHBG) which represents a soluble steroid hormone transporter protein (Grishkovskaya et al. 2002). This multifaceted interaction spectrum of LG domains led to their alternative naming “laminin-neurexin-SHBG” (LNS) domains (Missler and Südhof 1998).

ADGRG4 (former: GPR112) is an orphan aGPCR which was identified as a specific marker for enterochromaffin cells of the small intestine and neuroendocrine carcinoma cells (Leja et al. 2009). ADGRG4 is an evolutionarily old receptor being present in the genomes of all bony vertebrate classes (Wittlake et al. 2021). Among aGPCRs ADGRG4 is a poorly studied Gs protein-coupled receptor and limited information is available about its physiological relevance yet. Recently, the 7TM domain of ADGRG4 has been structurally characterized using cryo-electron microscopy (Xiao et al. 2022) displaying the same tethered *Stachel* peptide-mediated activation mechanism as for other aGPCRs (Liebscher et al. 2022). With more than 2,700 amino acid residues in its ECR, this receptor presents with one of the largest N-termini within the aGPCR class. However, only three domains are predicted by homology within this huge ECR – the PTX-like domain, the hormone receptor binding motif (HRM), and the GPCR autoproteolysis-inducing (GAIN) domain (Figure 1). The structural analysis of this ECR may provide clues of ADGRG4 function. GAIN and HRM domains of several other aGPCRs have been already structurally characterized by X-ray crystallography (Araç et al. 2012; Salzman et al. 2016; Leon et al. 2020). Because of its peculiar architecture as a highly conserved domain at the tip of a huge probably disordered region, we were interested in the structure of the PTX-like domain (ADGRG4-PTX) to understand its relevance in ADGRG4 function. Our X-ray crystallographic studies of ADGRG4-PTX revealed an overall fold similar to those of other PTX domains. However, ADGRG4-PTX homodimerizes Ca^2+^-independently, as also supported by SAXS measurements, and significantly differs in distinct structural features compared to known PTX structures. Our data strongly suggests that ADGRG4 may act as a mechanical force sensor for shear stress or changes in distance.

**Fig. 1.**
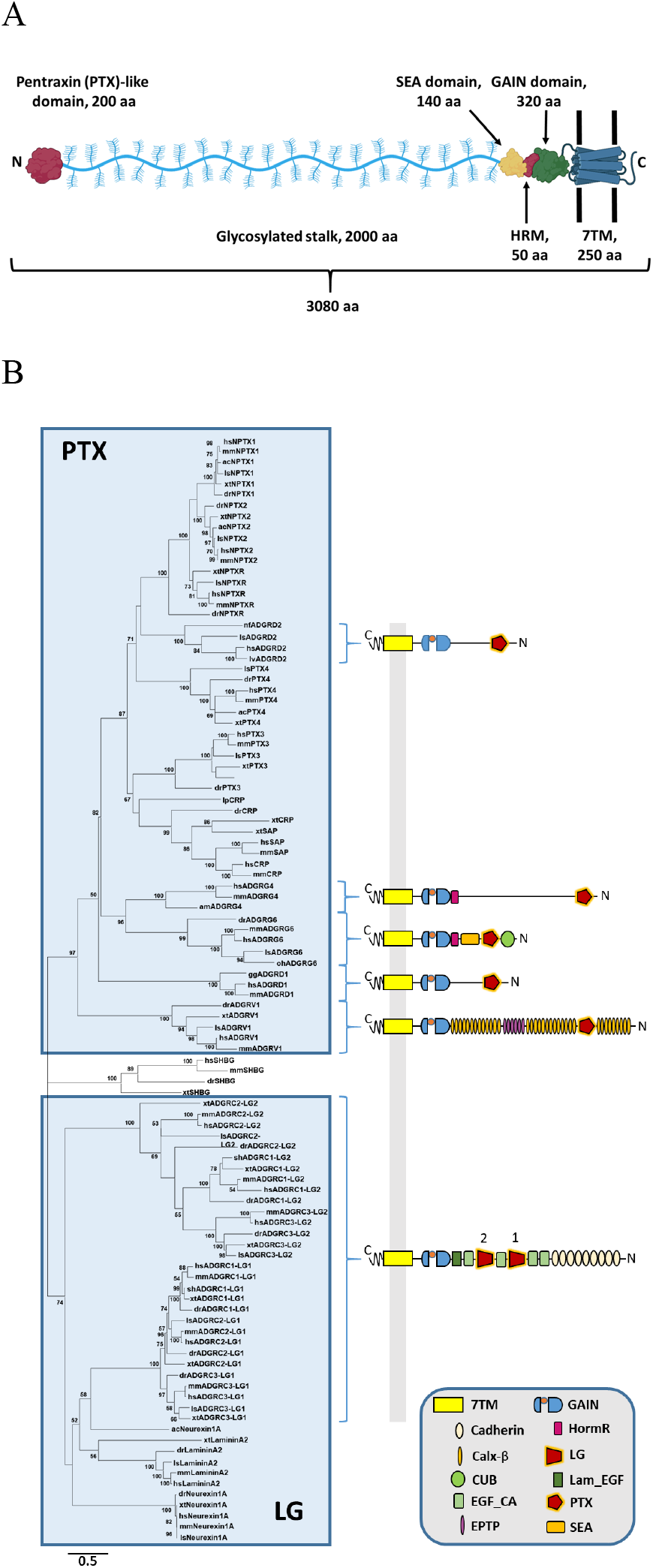
Predicted molecular architecture of ADGRG4 and phylogenetic relation of PTX-like domains. The structural organization of the human ADGRG4 is shown, predicted domains are indicated with their rough amino acid lengths. Created with BioRender.com A). Phylogenetic tree of selected PTX-like domains was generated by the maximum likelihood method (see Methods). Species ac: *Anolis carolinensis* (green anole), am: *Alligator mississipiensis* (American alligator), dr: *Danio rerio* (zebrafish), gg: *Gorilla gorilla*, hs: *Homo sapiens*, lp: *Limulus polyphemus* (Atlantic horseshoe crab), ls: *Lonchura striata domestica* (Bengalese finch), lv: *Lipotes vexillifer* (Chinese river dolphin), mm: *Mus musculus* (house mouse), nf: *Nothobranchius furzeri* (turquoise killifish), oh: *Ophiophagus hannah* (king cobra), sh: *Strigops habroptila* (kakapo), xt: *Xenopus tropicalis* (western clawed frog) B).

## Results and Discussion

### Phylogenetic analysis of aGPCR pentraxin-like and laminin-like domains reveals two distinct clusters

There are several PTX-like and LG domains in different soluble and membrane proteins. First, we analyzed their phylogenetic relations of the ADGRG4-PTX by aligning selected orthologous amino acid sequences of PTX-like/LG domains from different proteins and building a maximum likelihood tree. As shown in figure 1B, there are three main clusters as supported by bootstrap analysis – **cluster “PTX”** contains the PTX-like domains of the neuronal pentraxins NP1 and 2, the neuronal pentraxin receptor (NPR), CRP, SAP, PTX3, PTX4, as well as the aGPCRs ADGRD1, ADGRD2, ADGRG4, and ADGRG6. **Cluster “LG”** comprises the LG/PTX-like domains of neurexin 1α, laminin α LG2, and the LG1 and 2 domains of the aGPCRs ADGRC1–3. The third cluster lies in between the two other clusters and is exclusively formed by SHBG orthologs. Within the cluster **PTX**, the PTX-like domain of ADGRG4 clusters with those of ADGRG6 indicating close phylogenetic relation. However, the PTX-like domain of CRP and SAP seems to be more distantly related to the one of ADGRG4 which already indicates some significant structural differences to the prototypical domain. Interestingly, the two LG domains of the ADGRC1–3 independently cluster according to their position within the ECR, regardless of their paralog affiliation. This suggests that the two different LG repeats in ADGRC1–3 fulfill distinct functions.

### The N-terminus of ADGRG4 represents a conserved β-sandwich fold

For protein crystallization and structural analysis, the cDNA of the PTX-like domain of the human ADGRG4 (residues 1-240, UniProt accession number Q8IZF6-1) was cloned into a mammalian expression vector and transfected for secreted expression into the HEK293S GnTI^-^ cell line. The two-step purification (see Methods) yielded the protein in sufficient purity for crystallization (suppl. Fig. S1A). The protein exhibited a double band in a 12%-SDS-PAGE, which collapsed upon treatment with PNGase (suppl. Fig. S1B), indicating a heterogeneous glycosylation state. Crystals of space group C2 were obtained, which were optimized to a size of over 200 μm (suppl. Fig. S1C). Residues 26-162 und 176-231 are resolved in the electron density maps. The N-terminal residues 1-25 are the signal peptide and are not present in the secreted protein. The C-terminal residues as well as the loop residues 163-175 have no density, most likely due to flexibility. The crystallographic data revealed a β-sandwich fold (Fig. 2). Six and seven antiparallel β-strands assemble the two β-sheets A and B, respectively. The designation as “A” and “B” was adopted from the homologous PTX domains of CRP and SAP (Fig. 1B) (Ashton et al. 1997; Pepys 2018). The upper panel of figure 2 represents a top view onto the B face. This concave side of the β-sandwich features two protruding loops formed by residues 78-89 and 160-179 (Fig. 3), with the latter one being mostly unresolved in the electron density map. The A face is covered by another loop of residues 189-205 with a short α-helical stretch. This loop coordinates a metal ion via the side-chain carboxy group of the residue D202. This metal ion was modeled as Mg^2+^ based on an average coordination distance of 2.0 Å and the presence of 0.2 M MgCl2 in the crystallization medium (Fig. 2, suppl. Fig. S2A). The residual octahedral coordination sphere of the metal ion is completed by five water molecules, which in turn are not directly bound by other parts of the protein.

**Fig. 2.**
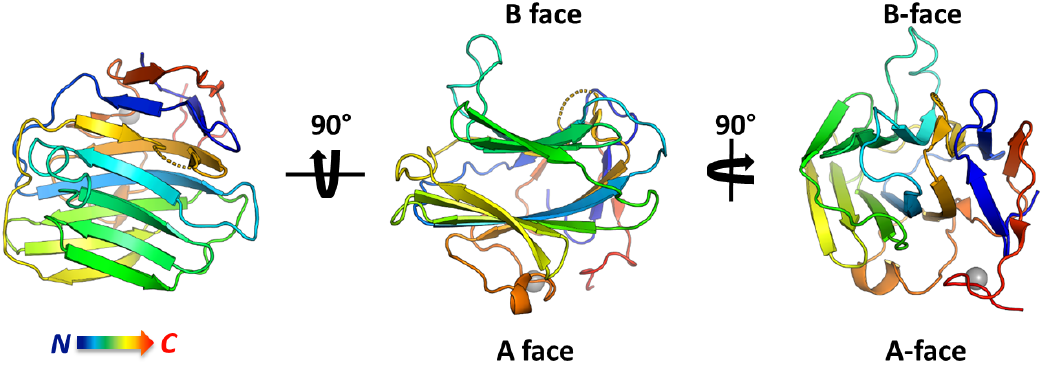
Tertiary structure of ADGRG4-PTX in a rainbow-colored cartoon representation. The grey sphere represents a bound Mg^2+^ ion.

**Fig. 3.**
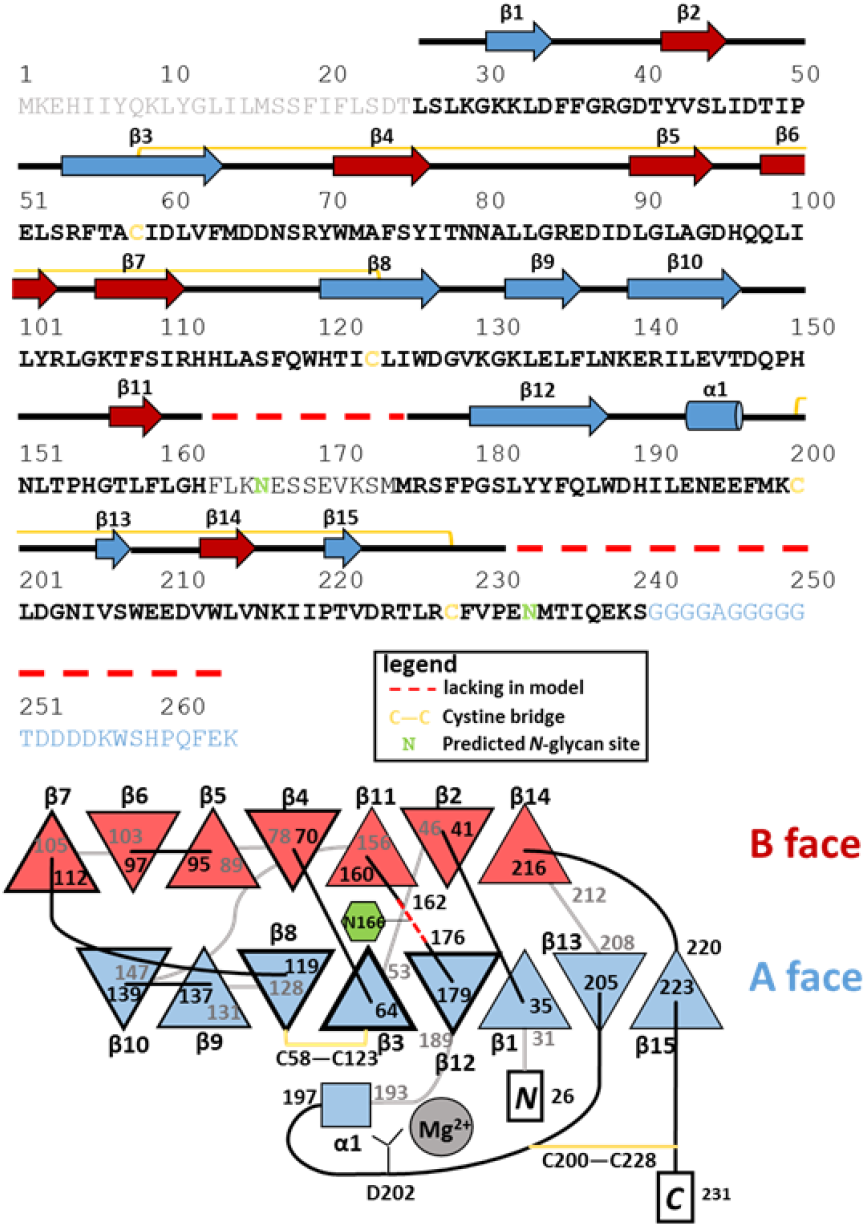
ADGRG4-PTX secondary structure (top) and topology diagram (bottom). Bottom scheme: Black/gray lines depict loops in front/rear of the panel plane. The nomenclature of the β-sheets as A face and B face is made according to the structural relatives CRP and SAP. Frame thickness of the β-strands correlates with their respective aa length. Note that this scheme is not drawn to scale.

Two disulfide bridges stabilize the protein fold (suppl. Fig. S2B). C58-C123 links strands β3 and β8 in the β-sheet of the A face (Fig. 3). C200-C228 connects the long A face loop 189-205 with the C-terminal residues of the PTX-like domain. This domain contains one putative N-glycosylation site at N166. The presence of N-glycosylation was verified by SDS-PAGE-based glycan staining (suppl. Fig. S1). Due to the disorder of residues 163-175, glycosylation at this site is not resolved in the crystal structure.

A multiple amino acid sequence alignment of 59 mammalian ADGRG4 orthologs was employed to determine the Shannon entropy as (Zheng et al. 2008) a measure of evolutionary conservation for each residue of the PTX domain (see supplements for species included). The result mapped onto the ADGRG4-PTX structure is shown in figure 4. Overall, there is a high degree of amino acid sequence conservation among the species, even for most surface residues. Especially the core strand residues of the β-sandwich exhibit the lowest Shannon entropy and thus highest conservation. The highest evolutional variability is found for the loop 64-70 connecting strands β3 and 4 and the unresolved loop 160-179. Longer stretches of extraordinary high conservation comprise the N-terminus including β1, β4, β8, and β9 including their connecting loop, β11 and β12. Interestingly, these conserved sections represent pairs of antiparallel β-strands: β1-β12, β8-β9 (A face), and β4-β11 (B face).

**Fig. 4.**
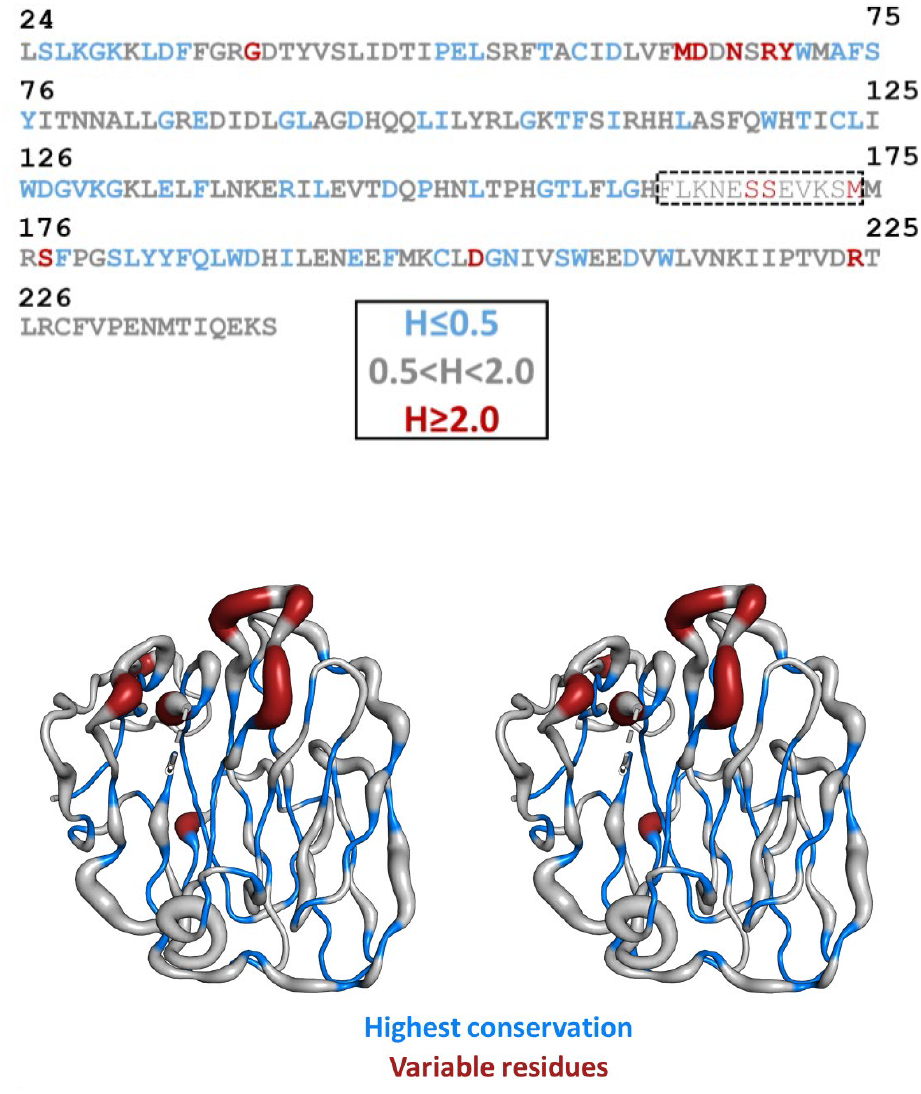
Shannon entropy based on multiple sequence alignment of 59 mammalian species plotted onto the structural model. View onto the ADGRG4-PTX B face in wall-eyed stereo display. The model shows the C_α_-trace in differing thickness. The thickness represents the Shannon entropy, H, of a given position and thus its evolutionary variability. Residues of extraordinary low variability are highlighted in blue, variable residues are shown in red. In general, a residue exhibiting H<2.0 is considered conserved (Perelson 1992; Garcia-Boronat et al. 2008). The unresolved loop is labeled in the sequence by a dashed box.

### The PTX-like fold exhibits distinct differences to structural homologs

To identify structural similarities the refined structural model of ADGRG4-PTX was submitted to the PDBeFold server for pairwise structural alignment with existing PDB structures (Krissinel and Henrick 2004). Based on the r.m.s.d. of the C_α_-chains, a selection of different structural relatives is given by Tab. S1. The closest structural relation is found for the human neuronal pentraxin 1 (NP1, 25 % identity) (Suzuki et al. 2020). The structures of the human CRP and human SAP follow very closely in this structural comparison. Figure 5 shows a multiple sequence alignment of the four proteins. The majority of the longer A face β-strands shows high conservation, with the B face being less conserved among the regarded pentraxins. Loops and outer β-strands are generally less conserved. Interestingly, the disulfide bridge C58-C123 (ADGRG4-PTX) that links two core β-strands of the A face is conserved among all structural relatives. However, the second disulfide bridge C200-C228 is not analogously found in CRP and SAP, but it is present in the NP1 structure. Furthermore, the amino acid sequence alignment indicates an ADGRG4-PTX section which does not align to the other sequences (marked yellow in Fig. 5). It forms the extended B face loop 53-66, which is not present in the related structures. The second B face loop 160-179 (marked blue in Fig. 5) is partially shared by all four pentraxins. However, the loop sequence of ADGRG4-PTX is mostly unrelated to its structural homologs and it is partially disordered, so the dashed line does not represent its actual length. In contrast, the other pentraxins share a high sequence identity and structural similarity in this area.

**Fig. 5.**
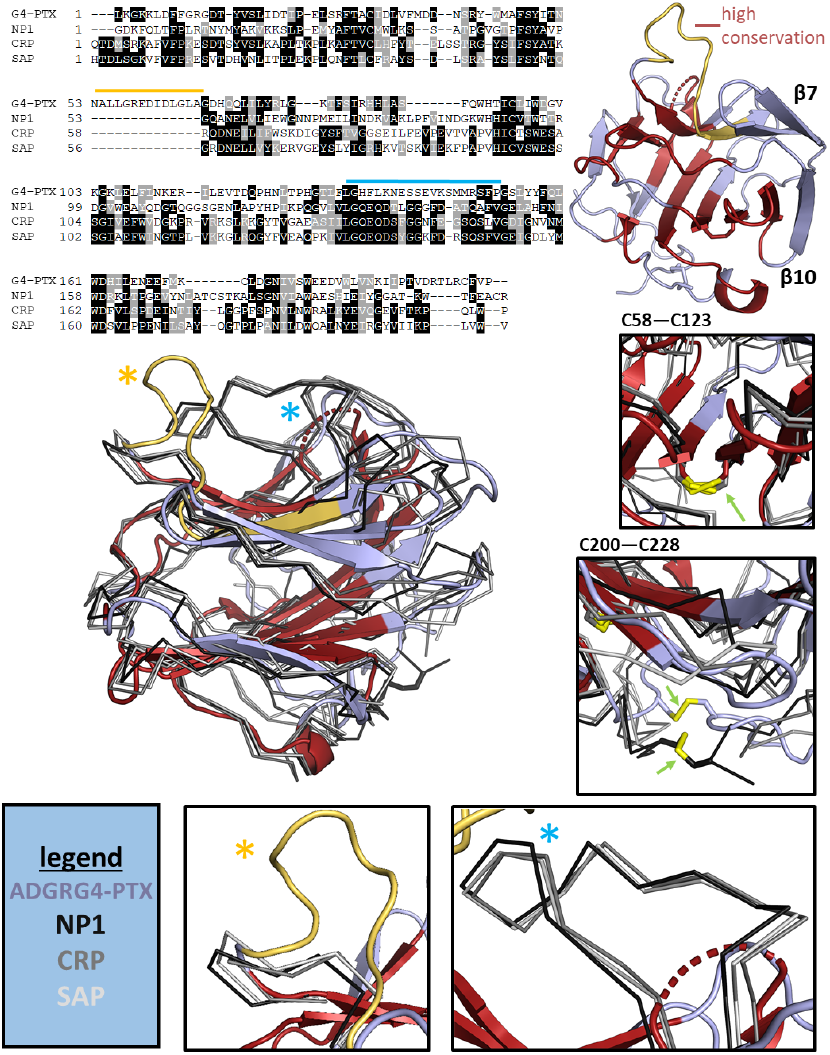
Alignment with closest structural homologs. Shown is the aa range of ADGRG4-PTX 26-231, NP1 (6ype) 225-429, CRP (1b09) 19-224 and SAP (2a3y) 20-223. Regions of the highest sequence consensus (based on the TCoffee server (Notredame et al. 2000)) are highlighted in red in the ADGRG4-PTX structure. Boxes showcase distinct features of ADGRG4-PTX in comparison to its structural relatives. Note that disulfide bridge C58-C123 is present in all four structures, whereas disulfide bridge C200-C228 is only found in ADGRG4-PTX and NP1. The box with the yellow asterisk shows an extended loop which is solely present in ADGRG4-PTX. The box with the blue asterisk depictures a close-up view on the long loop of the B face, which exists in all structures, but is unresolved (flexible) in the ADGRG4-PTX structure.

### The N-terminus of ADGRG4 shows Ca^2+^-independent homodimerization

Initial dynamic light scattering (DLS) of ADGRG4-PTX indicated oligomerization based on a hydrodynamic radius that was too large for an expected monomer mass of 30.5 kDa. To obtain more precise information about the molar mass in solution, static light scattering (SLS) was performed (suppl. Fig. S3). Indeed, the molecular mass determined by SLS increased with protein concentration and indicated a monomer-dimer-equilibrium in solution. The unit cell of the ADGRG4-PTX crystals contained a single protein chain as asymmetric unit. To address a potential homodimer interface, the respective symmetry mates in the C2 space group were assessed. Four different types of protein-protein contacts are present within the crystal lattice (suppl. Fig. S4). However, there is only one interface generated by point symmetry that exhibits a sufficient surface area for homodimerization. Figure 6 gives an overview of the interactions that constitute this dimer interface. The two protomers interact such that the two strands at the edge of the sandwiched β-sheet interact with those of the other protomer (β7 with β10’ and β10 with β7’, Fig. 3, Fig. 6). By this interaction two large continuous β-sheets are formed across the dimer interface. Core interactions are mediated by peptide backbone hydrogen bonds between each pair of opposing antiparallel strands (Fig. 6). In addition, the apolar side chains between the sandwiched β-strands 7 and 10 interact at the dimer interface via hydrophobic interactions and with tight steric fit. The peptide backbone hydrogen bonds are supported by several side-chain hydrogen bonds. Here, the tightest interaction is formed between the carboxamide-O_ε_ of Q98 from β-strand 6 (Fig. 6) and the guanidinium moiety of R141’ of β10. Additional hydrogen bonds are established between residues T107-T146, T107-E144’ (via a water molecule) and S109-E144’ (via a water molecule, Fig. 6). An ionic interaction is possible between R111 and E140’ at the interface flanks, however, for both residues the density indicates significant side-chain flexibility.

**Fig. 6.**
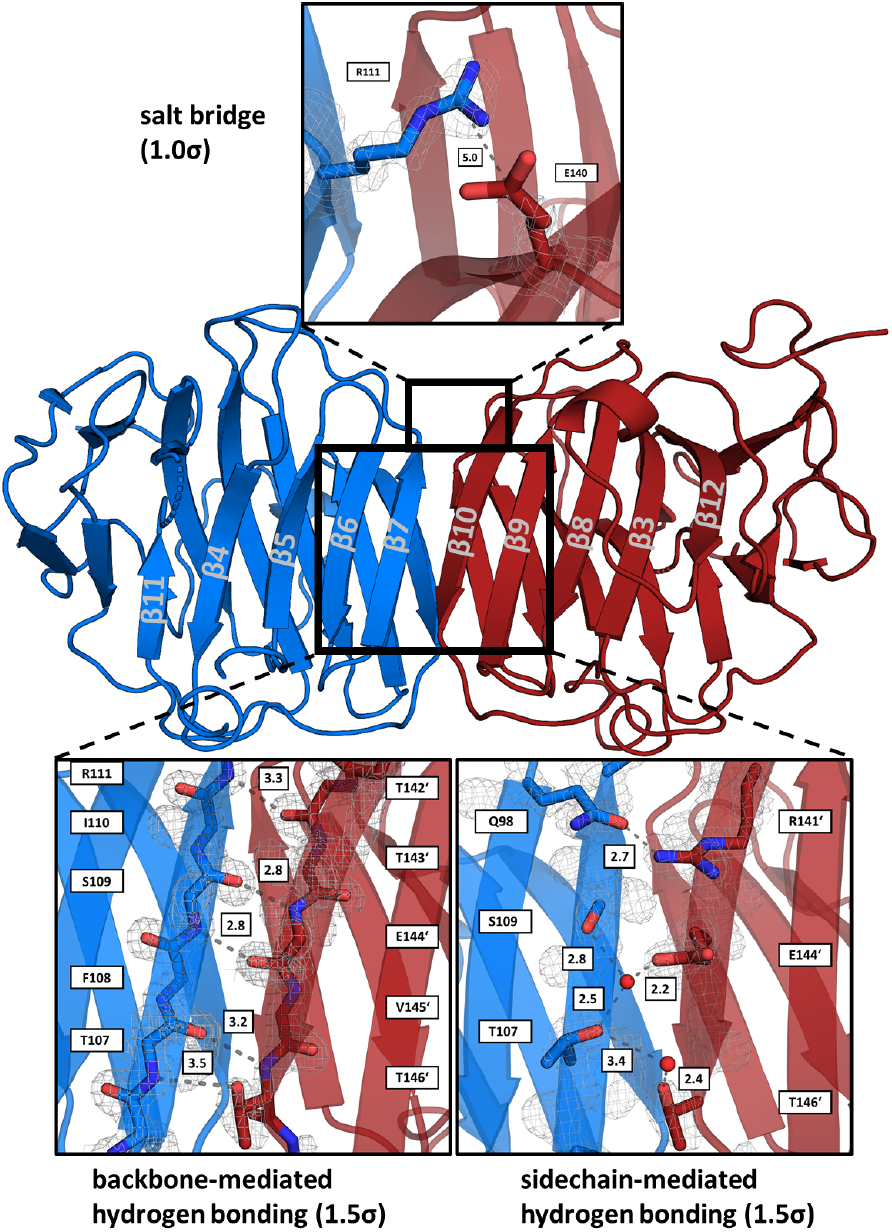
Interactions at the homodimer interface. For a better overview, backbone-mediated hydrogen bonds are shown separated from side chain-only interactions. Distances are stated in Å next to the dashed connectors. The respective contour levels of the (2Fo-Fc)-type electron density map are given next to the images.

To further characterize the monomer-dimer-equilibrium, SAXS experiments were performed at different AGDRG4-PTX concentrations. The scattering profiles exhibited concentration-dependent changes, which are highlighted by the observed R_g_ (radius of gyration) and D_max_ (maximum diameter of particle) values at different ADGRG4-PTX concentrations (suppl. Tab. S3). We determined the volume fractions of monomer and dimer for the scattering curves obtained at different protein concentrations via modeling with the program OLIGOMER (Fig. 7) (Konarev et al. 2003). Figure 8c shows a plot of the dimer volume fraction against c(ADGRG4-PTX). The data fit to the following binding model (with X_Dimer_ being the molar fraction of homodimer in the equilibrium):

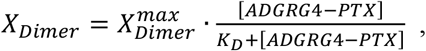

with a dissociation constant of K_D_ = (16±1) μM (Bisswanger 2017). This micromolar affinity characterizes the ADGRG4-PTX homodimer as a transient protein-protein interaction (Perkins et al. 2010). The discrepancy χ^2^ between fit and data increases at higher protein concentrations, which is due to larger deviations in the low q-region (q≤ 0.15 Å^-1^). This finding is probably due to additional intermolecular interactions at higher protein concentrations. The calculated scattering curves of the symmetrical dimer models 1 and 2 (Figure S4) were too similar to experimentally distinguish the two dimer models in the presence of the observed monomer-dimer mixture.

**Fig. 7.**
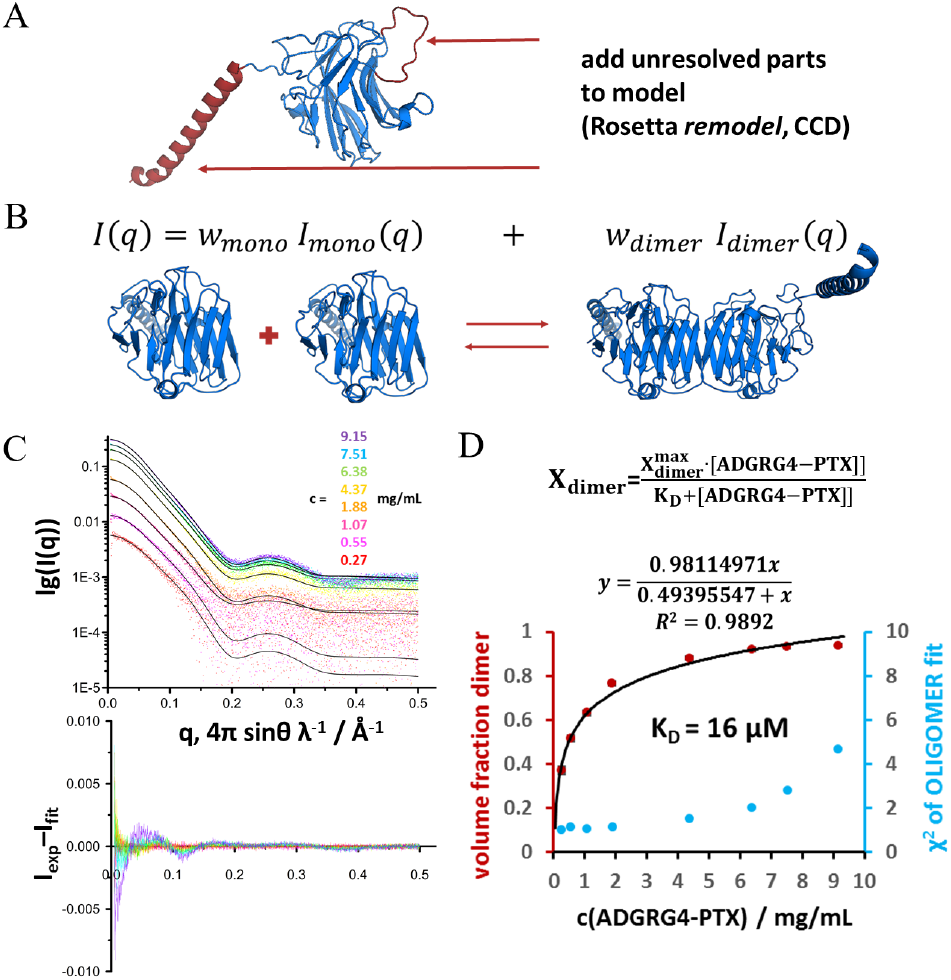
Overview of OLIGOMER modeling and results. To obtain a reasonable model for fitting to the SAXS data, the crystal structure was modified by adding the unresolved loop and C-terminus using Rosetta remodel (Leaver-Fay et al. 2011) A). Scheme of the principle of OLIGOMER (Konarev et al. 2003) B). Summary of the results obtained from OLIGOMER. On the left the scattering profiles are shown in semi-logarithmic manner with the respective modeled fit curves (black) C). The volume fraction of the dimer as calculated by the OLIGOMER fit is plotted against the protein concentration. A hyperbolic fit of the volume fraction vs. protein concentration yields a K_D_ of 16 μM for the homodimerization. X_dimer_-volume fraction of dimer, derived from OLIGOMER D).

**Fig. 8.**
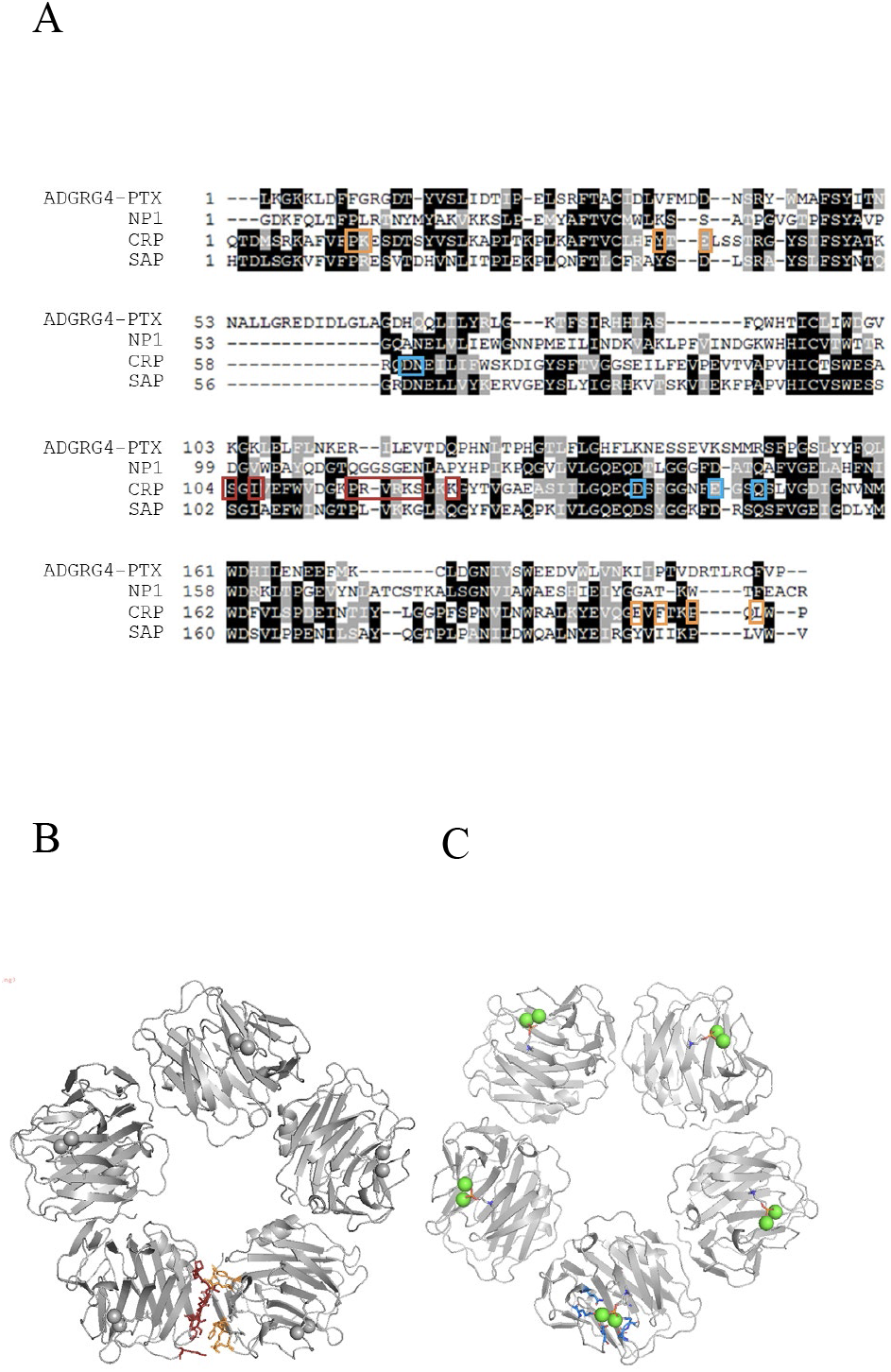
Alignment of ADGRG4-PTX with structural homologs A), highlighting CRP pentamer interfaces B) and Ca^2+^-binding sites C). Note, there is no sequence similarity between ADGRG4-PTX and CRP in the highlighted regions.

The classical pentraxins, CRP and SAP, feature characteristic Ca^2+^-binding sites, which are indispensable for ligand binding and, in case of CRP, pentamer stabilization (Fig. 8) (Pepys 2018). However, in the elucidated structure, only one bound metal ion was detected (see Fig. 2 and suppl. Fig. S2A), which is only loosely coordinated by a single carboxylate side chain. A potential influence of physiological Ca^2+^ concentration on the protein’s quaternary structure was assessed by DLS, SLS, nanoDSF, and crystallography. No significant change in the hydrodynamic radii and molecular mass was observed in the presence of Ca^2+^ (suppl. Fig. S5a). However, CaCl2 addition does increase the melting temperature Tm by 1.6 K, indicating a stabilizing effect on the ADGRG4-PTX fold (suppl. Fig. S5b). To study potential structural consequences of Ca^2+^ presence, crystals were soaked with 100 μM, 1 and 10 mM CaCl2 for 1 h or 19 h. Furthermore, the protein was also crystallized in presence of 10 mM CaCl2. At least one dataset per condition was recorded and analyzed. However, none of these datasets revealed new features in the electron density compared to the previous results without Ca^2+^ addition (data not shown). Support for a Ca^2+^-independent dimerization comes also from the comparison of the pentamer interfaces and the Ca^2+^-binding sites of CRP with the ADGRG4-PTX structure (Fig. 8). The pentamer-forming interaction sites in CRP and the corresponding positions in ADGRG4-PTX show low conservation (suppl. Fig. S6). This makes an ADGRG4-PTX oligomer formation similar to CRP unlikely. Furthermore, the Ca^2+^-coordinating carboxyl groups in CRP are not conserved in ADGRG4-PTX (Fig. 8). These facts together with the distant phylogenetic relation between CRP and ADGRG4-PTX (Fig. 1B) suggest that the PTX domain does not form Ca^2+^-dependent pentameric homophylic interactions.

As described before, the ADGRG4-PTX dimer interface is mainly formed by interactions between strands β7 und β10 at the edge of the central β-sheets. Since the hydrophobic side chains between these sandwiched β-strands are relatively well-conserved for their structural function in the densely packed core of the fold, it is difficult to deduce an additional functional role in dimer formation by conservation analysis (as opposed to generally less-conserved solvent exposed surface residues). We therefore analyzed the prediction of dimer structures by the AlphaFold machine learning algorithms (Jumper et al. 2021). AlphaFold uses two main sources of structural information for its prediction: Experimentally determined template structures of related proteins and coevolution of residues in spatial proximity in the folded protein. The first ranked dimer model matches the experimental ADGRG4-PTX dimer structure well (suppl. Fig. S7). Also, the model confidence of the dimer structure is high as demonstrated by the low predicted aligned errors between the protomers (PAE parameter in suppl. Fig. S7). As the template structures involved in modelling the dimer structure did not include structures that resemble the ADGRG4-PTX dimer, the model is predominantly based on information from sequence coevolution. Next, we also predicted the dimer structure for an additional 12 out of the 59 mammalian ADGRG4 homologs of the multiple sequence alignment (Fig. 4, suppl. Fig. S8). The sequences were chosen to evenly cover the phylogenetic tree. This analysis indicates that formation of this dimer is a conserved feature for all mammals and that it has functional relevance for the receptor (see Figure legend of suppl. Fig. S8 for further details).

### The putative dimer interface resembles that of SHBG and lectins

A very similar dimerization mode as found for ADGRG4-PTX has been described for the SHBG N-terminal LG domain (Grishkovskaya et al. 2002; Round et al. 2020). Here, the N-terminal LG domain provides both the steroid-binding site and the homodimer interface. However, this dimerization is Ca^2+^-dependent and facilitated by ligand binding (Bocchinfuso and Hammond 1994).

Recently, Leon et al. elucidated the structure of the zebrafish ADGRG6-ECR, which contains an internal PTX-like domain (Leon et al. 2020). However, the authors did not comment on the oligomeric state of the ECR in solution and the crystal structure does not feature a comparable dimer interface (PDB 6V55). As the ADGRG6-PTX domain represents the closest relative to ADGRG4-PTX, we also investigated ADGRG6 oligomerization. Therefore, the ECR of ADGRG6 was expressed in HEK293 cells, purified and analyzed by SEC-MALS-SAXS coupling (to be published). Conversely, the obtained molecular mass matches perfectly the expected one of a monomeric ECR. Furthermore, there is only poor sequence homology between ADGRG4-PTX and ADGRG6-PTX within the two strands β7 and β10 that form the putative homodimer interface (compare Figs. 3 and 9). AlphaFold predictions for the PTX-like domains of ADGRD1, ADGRD2 and ADGRG6 do not indicate dimer formation (suppl. Fig. S9). Hence, we conclude that our observations for ADGRG4-PTX are not generalizable for other PTX-like domains found in aGPCRs.

**Fig. 9.**
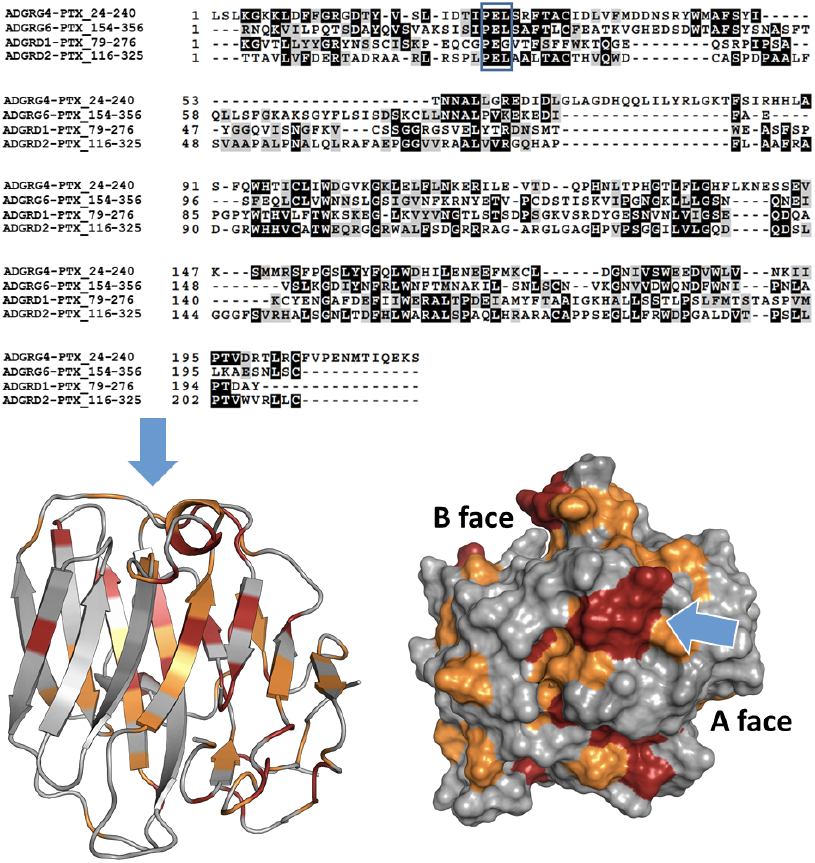
Alignment of ADGRG4-PTX with homologous PTX-domains of the human aGPCR members ADGRG6 (UniProt Accession Number: Q86SQ4-1), ADGRD1 (Q6QNK2-1), ADGRD2 (Q7Z7M1-1), a blue box highlights a conserved patch of surface residues. Bottom: ADGRG4-PTX (view onto B face) with highlighted residues of high (red) and medium (orange) sequence consensus. Blue arrows indicate a conserved patch at the flank of the structure with a core “PEL” motif.

### The typical binding rim of LG domains shows high surface conservation in ADGRG4-PTX

Despite their phylogenetic proximity, ADGRG4-PTX differs significantly from classical PTXs. This finding is supported by the non-PTX-like features of this “PTX-like” domain: There is no apparent Ca^2+^-dependent pentamerization. In contrast to classical PTXs, the functionally important B face does not exhibit the typical Ca^2+^-binding site (Fig. 8). Instead, the respective loop 160–179 is highly disordered and of comparably low evolutionary conservation with almost no sequence similarities to CRP or SAP (Fig. 5). In contrast to their high structural similarity, the main ligand-binding interface of PTX and LG domains differs significantly. While PTXs bind their ligands via the B face of the β-sandwich and its characteristic Ca^2+^-binding sites, LG domains employ the β-sandwich rim opposite to the N- and C-terminus as interaction area. This binding site is mainly established by the loops connecting strands β2–3 and β10–11. Diversity in these loops enables a broad ligand spectrum, with the core β-sandwich providing a rather rigid scaffold (Rudenko et al. 2001). In adhesive interactions between LG domains and proteoglycans, a divalent cation is typically involved which is only weakly coordinated by one or two residues of the LG domain. For example, the LG4 and 5 domains of the α-subunit of laminin-2 bind a glycan chain of α-dystroglycan mediated by bound Ca^2+^. A similar mechanism is proposed for the heparin proteoglycans agrin and perlecan (Dempsey et al. 2019). For the binding of steroid hormones to SHBG, the same β-sandwich rim area is essential again. In contrast to ADGRG4-PTX and other LG domains, in SHBG the loop linking strands β10 and β11 shows high disorder, which is probably critical for steroids to intercalate the β-sheets (Grishkovskaya et al. 2000). Intriguingly, an alignment of all PTX-like domains present in aGPCRs reveals a conserved patch of surface residues, located within the β2-3 loop around a “PEL” consensus sequence (Fig. 9). The central, highly conserved glutamate residue (E51 for ADGRG4) protrudes from the surface (suppl. Fig. S10). This finding suggests a common interaction interface for ADGR-PTX-like domains, which resembles that of the LG domain binding rim.

### Possible implication of the ADGRG4-PTX domain structure on *in vivo* functions

In principle ADGRG4-PTX homodimerization can occur at the same cell (*cis*) or between different cells in close proximity (*trans*) expressing the receptor protein. Homophilic interactions in *trans* were previously shown for other aGPCRs. For example, ADGRC1 (CELSR1) undergoes homophilic *trans* interactions and concentrates at cell-cell contacts in adherens junctions, where it is involved in the planar-cell-polarity (PCP) pathway required for neural plate bending (Nishimura et al. 2012). *Trans* dimerization was found also for the homologs ADGRC2 and 3 (Shima et al. 2007). Similarly, Flamingo, a structurally related aGPCR in the fly *D. melanogaster*, is localized at cell-cell boundaries of Drosophila wing cells and engages in PCP signaling (Usui et al. 1999). In both cases, *trans* homodimer formation is mediated by repeats of cadherin domains within the ECR. ADGRG1 (GPR56) represents another example of a homophilic *trans* interaction of the ADGRG subfamily. Here, Paavola et al. showed that the homophilic N-terminal interaction enhances ADGRG1-mediated RhoA activation (Paavola et al. 2011).

Given the size of over 2,700 amino acid residues of the ADGRG4 ECR, which is even larger than the ECRs of ADGRC, it is reasonable to assume homophilic *trans* interaction mediated by the N-terminal PTX-like domain. The highly glycosylated mucin-like stalk (Fredriksson et al. 2002) between the PTX-like and HRM domain might act as a spacer to span intercellular clefts, enabling the ADGRG4-PTX to reach remote binding partners. An unfolded protein has a contour length of approximately 4 Å per residue (Ainavarapu et al. 2007), but the chain is highly flexible and forms mostly globular random coil structures. It has been shown for mucins, that the O-glycosylation increases the stiffness (persistence length) of the peptide core about 10-fold resulting in the formation of an extended structure, such that the radius of gyration increases almost 10-fold (Gerken et al. 1989). With a pitch of 2.5 Å, as determined for mucins, a fully extended ECR-chain of ADGRG4 may span a distance of more than 0.5 μm. Recently, Malaker *et al*. developed an algorithm to detect mucin-like domains within the human proteome (Malaker et al. 2022). The algorithm first identifies O-glycosylation sites via the NetOGlyc 4.0 server (Steentoft et al. 2013). Overlapping phosphorylation sites are subtracted from the total O-glycosylation sites as false positives. Next, based on four mucin-typical benchmarks, the algorithm determines a so-called “mucin score”. A mucin score of >2 represents a high confidence for the mucin domain assignment. Values between 1.5–1.2 represent a low confidence, while lower scores are not considered mucin-like. Analysis of ADGRG4 results in a mucin score of 3.4, which clearly indicates the mucin-like properties of the receptor. The algorithm detected a total of 212 mucin-domains, which arrange seamlessly starting at residue 279 until residue 2244 (Uniprot Q8IZF6). This region starts C-terminally to the PTX-like domain and ends roughly 140 amino acids N-terminal to the predicted HRM. Interestingly, ColabFold (Mirdita et al. 2021) predicted this ~140 amino acid stretch to establish a Sperm protein, Enterokinase and Agrin (SEA) domain (suppl. Fig. S11). SEA domains are a typical feature of many mucins: MUC1, MUC3, MUC12, MUC13 and MUC17 mucins all possess a single, and MUC16 even multiple SEA domains (Pelaseyed et al. 2013). However, the typical autoproteolytic site G↓S[V/I]VV present in many mucin SEA domains is lacking for the ADGRG4 SEA domain. The presence of SEA domains has been reported already for three other aGPCR ECRs, namely ADGRF1 (GPR110), ADGRF5 (GPR116) (Fredriksson et al. 2002; Lum et al. 2010) and ADGRG6 (GPR126) (Leon et al. 2020). The homodimerization of ADGRG4’s N-terminal domain presented in this study strongly suggests a homophilic *cis* but also a *trans* dimerization (<1 μM distance) of this receptor. We determined a K_D_ value of 16 μM for the dimerization *in vitro* (Fig 7C). This translates into a binding ΔG of 28.5 kJmol^-1^ at 37 °C. Interestingly, the dimer interface does not mask the typical LG domain binding rim. Therefore, it is likely that additional ligands may bind to the ADGRG4-PTX homodimer.

ADGRG4 has been discovered as a specific marker of enterochromaffin (EC) cells (Leja et al. 2009) and Paneth cells (Franzén et al. 2019). EC cells are responsible for serotonin production and secretion and are most abundant in the mucosa of the duodenum (Ito et al. 2009). The mucosa of the small intestine forms a surface of villi and crypts, which contributes to a large overall surface area gain. The crypts represent deep but very narrow clefts. Here, EC cells are mainly localized (GRAEME-COOK 2009). EC cells feature a luminal side which forms microvilli, as well as a basolateral border that is in contact with afferent and efferent nerve terminals located at the lamina propria (Bistoletti et al. 2020). Similarly, Paneth cells are specialized secretory epithelial cells mainly localized in the crypts of the small intestine. Given the large ECR of ADGRG4 which is potentially able to span a distance of close to 0.5 μm, homophilic *trans* interactions between cells at opposite sides of the crypt lumen may represent a plausible scenario. Together with the general notion of aGPCRs to act as mechanosensors (Petersen et al. 2015; Wilde et al. 2016; Scholz et al. 2017; Yeung et al. 2020) we hypothesize that ADGRG4 acts as a distance sensor within epithelial crypts of the small intestine via *trans* homodimerization (Fig. 11). Movements of the intestinal tract may lead to transient dilations of the crypts. Alternatively, epithelial layer growth may separate *trans*-interacting cells. Once a maximal distance of ~1 μm is reached (considering two ADGRG4 *trans* dimer molecules), traction forces are generated due to the homodimer interaction. These forces could be transmitted via the stalk region to the GAIN domain, resulting in an exposure of the agonistic *Stachel* sequence or activity-relevant isomerization of the tethered agonistic sequence (Fig. 10) (Liebscher and Schöneberg 2016; Schöneberg et al. 2016). To investigate whether such *trans* interaction is feasible we localized ADGRG4 in small intestine with different methods. First, we applied an anti ADGRG4 antibody directed against the ECD of the receptor. As shown in Figure 11A, immunopositive cells were mainly scattered in the intestinal epithelium of crypts and villi. Next, a specific probe to detect ADGRG4 mRNA was used and epithelium cells were co-stained with an anti-β-catenin antibody (Fig. 11B). Again, ADGRG4-expressing cells were mainly scattered in the intestinal epithelium. However, we did not find opposed ADGRG4-expressing cells rejecting the hypothesis of a *trans* homodimerization as the major form of interaction. Therefore, the *cis* homodimerization is more likely. Together with the fact that the highly glycosylated mucin-like stalk between the PTX-like and HRM domain has an increased stiffness and is most likely orientated to the intestine lumen, we propose that ADGRG4 acts as a mechanosensor for shear forces.

**Fig. 10.**
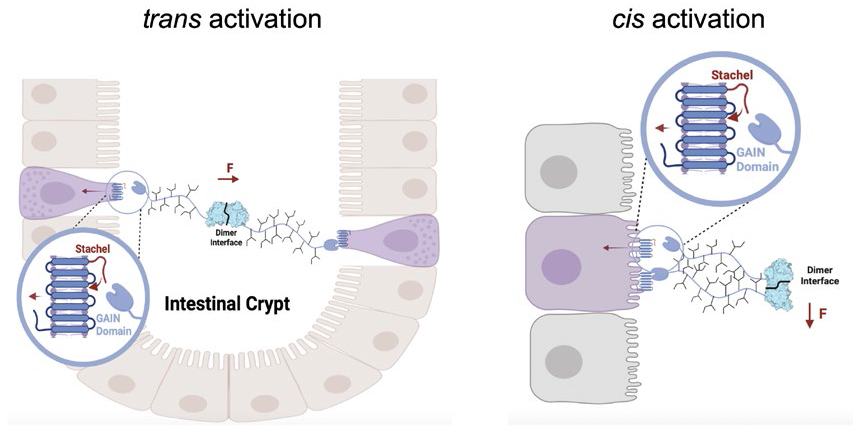
Scheme of hypothetic homophilic *trans* and *cis* interaction of ADGRG4 in an intestinal crypt. AGDRG4 may form *trans* dimers via homophilic PTX interaction. Upon bowl movements or cell proliferation-mediated changes in the cell-cell distances forces (F = force) may emerge at the homodimer that are conducted to the GAIN domain (blue-colored domain) via the stalk. This leads to the exposition of the *Stachel* to activate the 7TM. Black branches at the stalk indicate its high amount of glycosylation. In a *cis* interaction between ADGRG4 expressed in the same cell, shear forces my lead to receptor activation. Created with BioRender.com.

**Fig. 11.**
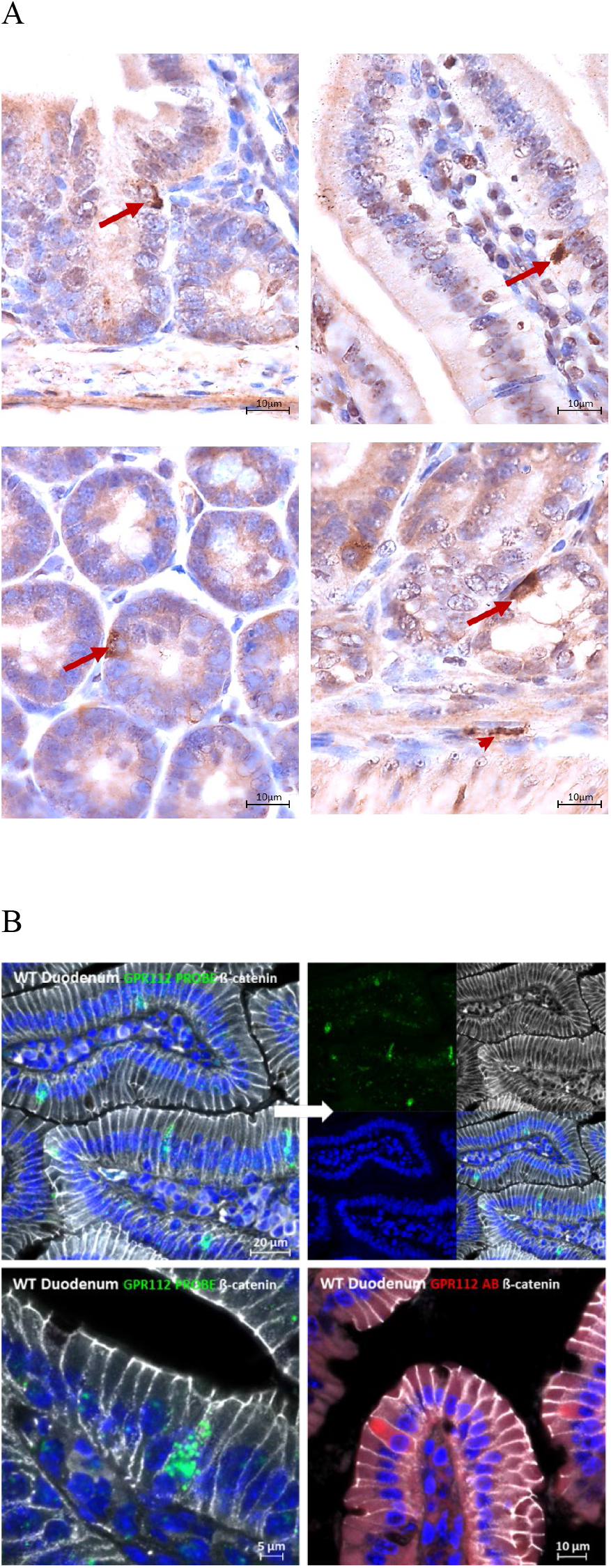
Detection of ADGRG4 protein and m-RNA in small intestinal tissue. Immunohistochemical detection of ADGRG4 positive cells in small intestinal epithelium (arrows) and underlying layers (arrowhead) A). Detection of ADGRG4-mRNA (green, FISH) or ADGRG4-protein (red, IF) positive cells in small intestinal epithelium highlighted by IF-ß-catenin (white) and DAPI-nuclear staining (blue) B).

In sum, our findings show a specific PTX structure enabling ADGRG4 to homodimerize via this domain. Reflecting the unique structure of ADGRG4, the activation mechanism by exposing a tethered sequence in its active conformation (*Stachel*) to the 7TM and its highly cell-specific expression one can assume that this aGPCR is a shear stress mechanosensor in EC and Paneth cells of the small intestine.

## Experimental Procedures

### Materials

All chemicals, reagents and compounds used in this study were purchased from AppliChem (Darmstadt, Germany), Carl Roth (Karlsruhe, Germany), Sigma-Aldrich (Steinheim, Germany), SERVA (Heidelberg, Germany), and Roche (Basel, Switzerland). Cell culture media and supplement were obtained from Gibco®, Life Technologies™, Thermo Fisher Scientific (Waltham, MA, USA). Enzymatic reagents and buffers were purchased from New England Biolabs (Frankfurt a. M., Germany), Thermo Fischer Scientific and Fermentas (St. Leon-Rot, Germany). Molecular biology kits were ordered from Qiagen (Hilden, Germany). Prepacked columns for protein purification were purchased from GE Healthcare (Buckinghamshire, UK). Primers were obtained from biomers.net GmbH (Ulm, Germany). To verify cloning results, Sanger sequencing was performed by Eurofins Genomics Germany GmbH (Ebersberg, Germany).

### Methods

#### Preparation of ADGRG4-PTX-like domain

##### Molecular cloning

The cDNA encoding for the N-terminus of the human ADGRG4 was synthesized by the Thermo Fisher GeneArt service. For subcloning into the pVitro2-EGFP backbone, the following primers were ordered: forward: 5′-TATA<u>ACGCGT</u>GCCACCATGAAGGAACACATCAT-3′ (*Mlu*I), reverse: 5′-GTAC<u>GTCGAC</u>TCATTATTTCTCGAACTGGGGGTG-3′ (*Sal*I). The final expression vector comprised the ADGRG4 amino acid sequence 1-240 (referred to as ADGRG4-PTX) with a GS-linker, enteropeptidase site and StrepII-tag at the C-terminus (suppl. Fig. S1). The vector backbone also contained a cassette for constitutive EGFP coexpression and a hygromycin resistance as selection marker for stable recombinant mammalian expression.

##### Generation of a stable cell line for large scale ADGRG4-PTX expression

A stable HEK293S GnTI^-^ cell line expressing the ADGRG4-PTX construct described above was established using hygromycin B resistance. The cell line was cultured in full medium, consisting of DMEM supplemented with 10% (v/v) fetal bovine serum, 1% (v/v) GlutaMax, 1% (v/v) non-essential amino acids and 1% (v/v) penicillin/streptomycin. For large scale expression, aliquots of the cryo-stored cell line were seeded of a single T-75 tissue culture flask and further expanded in T-175 (25×10^6^ cells). After reaching confluency, the cell were transferred into roller bottles (2125 cm^2^ surface area). The cell line was kept in roller bottles for two weeks, with two medium exchanges using reduced medium, which contained only 2% (v/v) of fetal bovine serum. The conditioned media of all passaging steps were stored at 4 °C and pooled. 1 mL of BioLock biotin blocking solution (IBA Lifesciences) were added to 1.5 L conditioned medium. The medium was gently stirred for 45-60 min at 4 °C and subsequently submitted to centrifugation at 16,000x g for 30 min. After that, the supernatant was filtered through a 0.22 μm polyether sulfone membrane (Techno Plastic Products). Prior to affinity chromatography, the filtered conditioned medium was tenfold concentrated by an ultrafiltration unit (10 kDa molecular weight cut-off (MWCO), GE Healthcare).

##### Purification

All chromatography steps were performed at 4 °C using the ÄKTAxpress and ÄKTA pure protein purification systems (GE Healthcare). For affinity purification of the ADGRG4 construct, three 5 mL StrepTrap HP (GE Healthcare) were connected to increase the amount of isolated protein per flow through. The washing buffer contained 100 mM Tris pH 8.0, 150 mM NaCl. The elution buffer additionally contained 2.5 mM D-desthiobiotin. After affinity chromatography, the preparation was further polished via size-exclusion chromatography (SEC). Here, a HiLoad^®^ 16/60 Superdex^®^ pg 200 (GE Healthcare) was applied for preparative runs. The eluate of the affinity purification was concentrated to a volume between 1-2 mL before injection using centrifugal filter units (Amicon, Merck millipore) with a MWCO of 5,000 Da. The SEC buffer contained 20 mM Tris pH 8.0, 150 mM NaCl.

##### Gel-based analyses

SDS-PAGE analysis was conducted after Laemmli. To stain protein glycosylations on SDS-PAGE gels, the periodic acid-Schiff-base method was employed (McGuckin and McKenzie 1958; Zacharius et al. 1969). Western blot analysis was performed via semidry electroblot onto a PVDF-membrane. After blocking the membrane with 3% (w/v) albumin in PBS-T over night, it was treated for 1.5 h with a StrepTactin horse radish peroxidase conjugate (IBA Lifesciences), diluted 1:75,000 in PBS-T. The membrane was washed twice in PBS-T and three times in PBS and readout was enabled using the ECL Select Western Blotting Detection Reagent (GE Healthcare).

#### Dynamic and static light scattering

Light scattering measurements were performed using a DynaPro® NanoStar® device (Wyatt Technologies). The sample was centrifuged prior to measurements for 10 min at 21,000x g. 3 μL of sample were pipetted into a quartz cuvette. After inserting the cuvette into the device, the system was allowed to equilibrate at 20 °C for 4 min. The integration time for autocorrelation was set to 5 s. Using the software DYNAMICS® (Wyatt Technologies), the autocorrelation function was analyzed by fitting it both in a cumulant and regularized manner. For SLS, a matching buffer blank was measured additionally.

#### X-ray crystallography

##### Crystallization

Crystals suitable for X-ray crystallography were obtained by hanging drop vapor diffusion by mixing 3 μL of protein solution at 3.1 mg/mL with 1 μL of reservoir solution containing 0.1 M Tris pH 8.5, 30 % polyethylene glycol (PEG) 4000 and 0.2 M MgCl_2_. Crystals between 100-200 μm length grew within a week.

##### Data collection and structure determination

Prior to X-ray diffraction experiments, suitable single crystals (min. 50-100 μm length) were transferred for a few seconds to a solution containing the respective crystallization reservoir composition and additional 20% (v/v) glycerol as cryoprotectant. Then, the crystals were snap-frozen in liquid nitrogen on nylon loops connected to goniometer-compatible magnetic mounts and stored in liquid nitrogen-immersed vials until measurement. Crystallographic measurements were performed using synchrotron radiation at beamline P14, operated by EMBL Hamburg at the PETRA III storage ring (DESY, Hamburg, Germany). Data were collected at λ=2.0664 Å (for phasing via anomalous dispersion, data not shown) or 0.9762 Å (Tab. S2) by an EIGER 16M detector. During data collection, the crystal was constantly cooled to 100 K via a nitrogen jet. Data reduction was performed by XDS (Sparta et al. 2016). Phasing was achieved by combining molecular replacement (MR) and sulfur single-wavelength anomalous diffraction (S-SAD) using PHASER and PHASER-EP (McCoy et al. 2007). The initial models were iteratively improved by refinement using phenix.refine (Liebschner et al. 2019) and manual rebuilding in COOT (Emsley et al. 2010) (suppl. Tab. S2). Molecular figures were generated with Pymol (www.pymol.org).

#### Small-angle X-ray scattering

SAXS measurements were performed using synchrotron radiation at beamline P12, operated by EMBL Hamburg at the PETRA III storage ring (DESY, Hamburg, Germany) (Blanchet et al. 2015). Data were collected at 1.23987 Å using a Pilatus 6M detector at 3.0 m sample-detector distance. The sample cell consisted of a horizontal thermostated (278-323 K) quartz capillary with 50 μm thick walls and a path length of 1.7 mm. The samples (30 μL) continuously flowed through the quartz capillary to reduce X-ray radiation damage. To minimize aggregation due to radiation damage, 3–5 % (v/v) glycerol was added to the samples.

The SAXS data were automatically processed on-site by the SASFLOW suite (Franke et al. 2012). Processing involves radial averaging of the two-dimensional scattering pattern, normalization against the transmitted beam intensity, detection and exclusion of frames suffering from radiation damage and buffer background subtraction.

#### Phylogenetic analysis

##### Cluster analysis

The sequences of PTX-like domains were taken from NCBI and aligned using MUSCLE implemented in MEGA11 (Stecher et al. 2020; Tamura et al. 2021). The evolutionary history was inferred by using the Maximum Likelihood method and JTT matrix-based model (Jones et al. 1992). The bootstrap consensus tree inferred from 1000 replicates (Felsenstein 1985) is taken to represent the evolutionary history of the taxa analyzed. Branches corresponding to partitions reproduced in less than 50% bootstrap replicates are collapsed. The percentage of replicate trees in which the associated taxa clustered together in the bootstrap test 1000 replicates) are shown next to the branches (ibid.). Initial tree(s) for the heuristic search were obtained automatically by applying Neighbor-Join and BioNJ algorithms to a matrix of pairwise distances estimated using the JTT model, and then selecting the topology with superior log likelihood value. This analysis involved 102 amino acid sequences. There were a total of 350 positions in the final dataset. Phylogenetic analyses were conducted in MEGA11 (Stecher et al. 2020; Tamura et al. 2021).

##### Shannon-entropy conservation scores

A multiple sequence alignment (MSA) of 59 mammalian ADGRG4 orthologs was generated using MUSCLE. Based on this MSA, the Shannon entropy of each aa residue was determined as a measure of evolutionary conservation (Shannon 1948). Subsequently, these Shannon entropy scores were plotted onto the crystal structure, employing the Protein Variability Server (Garcia-Boronat et al. 2008).

#### Structure prediction

The ColabFold (Mirdita et al. 2021) implementation of AlphaFold algorithms (Jumper et al. 2021) was used to predict dimer structures. Calculations were run on the ColabFold notebook AlphaFold2.ipynb. Multiple sequence alignments were generated via MMseqs2 (Mirdita et al. 2021) using sequences from UniRef. AlphaFold2-multimer was used for structure prediction and the “paired+unpaired” mode (ibid.) was chosen for complex prediction.

#### Cell localization experiments of ADGRG4 (GPR112) in murine duodenum

##### Immunohistochemical (IHC) and immunofluorescence (IF) staining

Small intestine was harvested from mice and fixed overnight at 4 °C in 4% phosphate-buffered formalin. After thorough rinsing with tap water, the tissue was dehydrated in a series of graded alcohols, passed through xylene, and embedded in paraffin vax. The embedded tissue was cut into 10 μm thick sections, deparaffinized in xylene and rehydrated in a series of graded alcohols. Prior to IHC-anti-GPR112-staining, hydrated sections were heated in sodium citrate buffer (pH 6.0, 10 min) to retrieve antigens, immersed in 3% H_2_O_2_ in phosphate buffered saline (PBS, pH7.4, 10 min, room temperature [RT]) to quench endogenous peroxidase activity, and incubated with 5% normal goat serum (30 min, RT) to reduce background staining due to nonspecific interactions between the secondary biotinylated goat anti-rabbit antibody and the section surface. Primary anti-GPR112 polyclonal antibody (cpa3095, Cohesion Biosciences, London, UK) was diluted 1:100 in antibody buffer (PBS, pH 7.4 containing 0.5% bovine serum albumin (BSA) and 0.3% Triton X-100) and applied overnight at 4 °C in a humidified chamber. Immune complexes were visualized by successive incubations with biotin-conjugated goat anti-rabbit secondary antibody (1 h, RT), avidin-biotin-peroxidase complex (Vectastain, Burlingame, CA, 30 min, RT), and a mixture of 3,3′-diaminobenzidine and urea-hydrogen peroxide (SigmaFast DAB tablets, Sigma-Aldrich, St Louis, MO, 30-60 sec, RT). The peroxidase/DAB reaction was stopped by rinsing the sections with 0.05M Tris-HCl buffer, pH 7.4. Finally, immunohistochemically labeled sections were counterstained with Mayer’s hemalum, dehydrated, and embedded with Roti-Histokit (Carl Roth, Karlsruhe, Germany). For immunofluorescence labeling, a 1:100 dilution of primary polyclonal anti-GPR112 antibody was applied together with primary monoclonal anti-ß-catenin antibody (1:100, #610154, BD Biosciences, Europe) on antigen-retrieved and background reduced sections overnight at 4 °C. Immune complexes were detected with a mixture of species-matched Alexa-568-and 647-conjugated secondary antibodies (1:400, Invitrogen, Carlsbad, CA, USA), whereas nuclei were counterstained with DAPI (4′,6-diamidino-2-phenylindole, Serva, Heidelberg, Germany). Finally, sections were embedded in fluorescent embedding medium from Dako (Aligent, Frankfurt, Germany). Fluorescence images were captured using an LSM700 confocal laser scanning microscope (Carl Zeiss AG, Jena, Germany).

##### *Fluorescence in situ* hybridization (FISH)

The RNAscope Multiplex Fluorescent Reagent Kit v2 (Advanced Cell Diagnostics [ACD], Berlin, Germany) was used to localize ADGRG4 mRNA on small intestinal sections. Tissue was fixed with 4% phosphate buffered formalin for 24 h and embedded in paraffin wax. Sections of 10 μm were made from the paraffin embedded tissue samples and mounted on Superfrost® slides. ADGRG4 RNA labeling was performed using a commercially purchased probe from ACD according to the manufacturer’s instructions. The 1:50 diluted probe was incubated on the tissue sections for 2 h at 40 °C in a manual HybEZ™ II assay hybridization system (ACD). Probe remaining on the tissue sections was amplified and labeled with a single Opal 520 fluorophore at a dilution of 1:750 (OP-001001, Akoya Biosciences, Marlborough). After RNA labeling, combined ß-catenin Alexa 647-IF staining and DAPI nuclear counterstaining was performed as described above. The now FISH-IF-stained sections were embedded and analyzed under the LSM 700 confocal laser scanning microscope (see above).

