## Supplements for "The dimerized pentraxin-like domain of the adhesion G protein-coupled receptor 112 (ADGRG4) suggests function in sensing mechanical forces"

*Correspondences should be addressed to:

MKEHIIYQKLYGLILMSSFIFLSDTLSLKGKKLDFFGRGDTYVSLIDTIPELSRFTACIDLVFMDDNSRYWMAFSYITNNALLGREDIDLGLAGDHQQLILYRLGKTFSIRHHLASFQWHTICLIWDGVKGKLELFLNKERILEVTDQPHNLTPHGTLFLGHFLKNESSEVKSMMRSFPGSLYYFQLWDHILENEEFMKCLDGNIVSWEEDVWLVNKIIPTVDRTLRCFVPENMTIQEKSGGGGAGGGGGTDDDDKWSHPQFEK*

Fig. S1: Sequence of the ADGRG4-PTX construct. Grey: ADGRG4 signal peptide; blue: ADGRG4 N-terminus (26-240); yellow: linker; green: enteropeptidase cleavage site; red: StrepII-tag

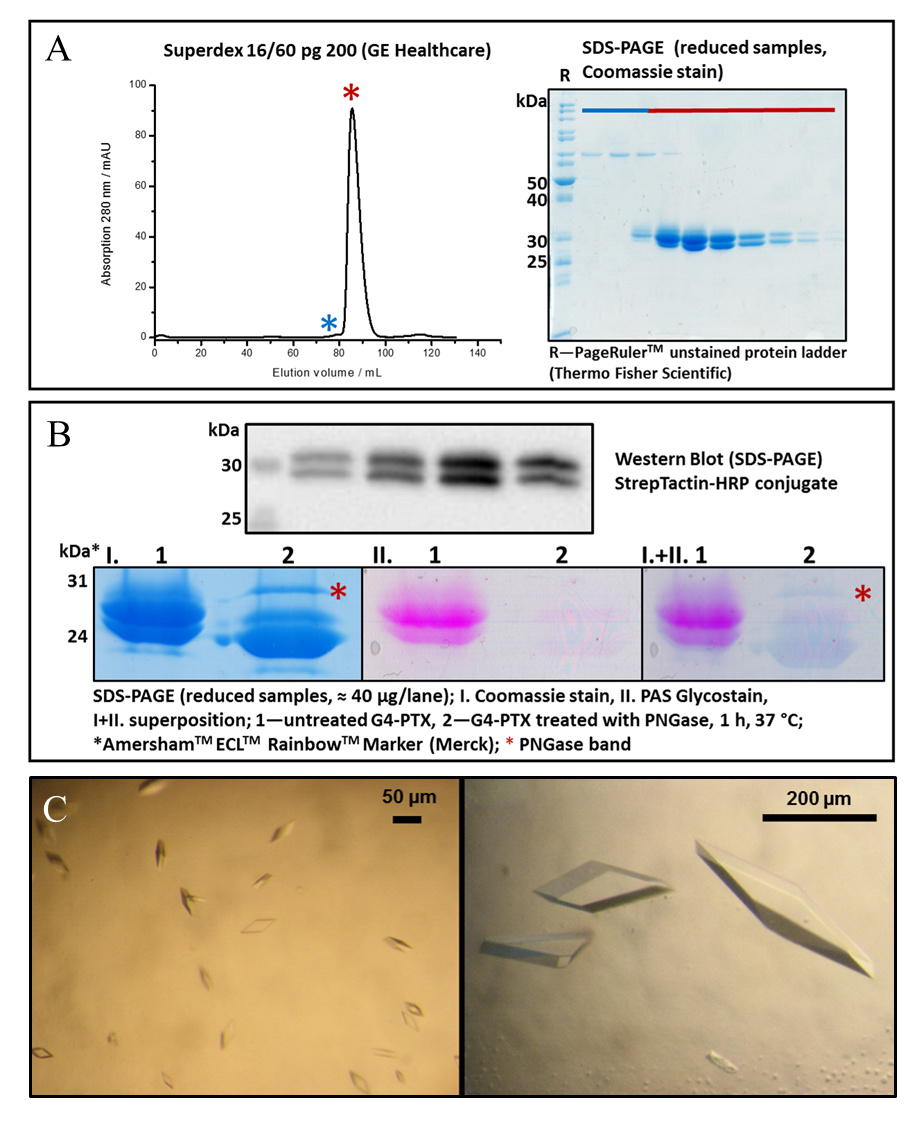

Fig. S1: Representative SEC chromatogram and corresponding SDS-PAGE (T = 12%) A). Band pattern of AGPRG4-PTX in 12 %-PA-gels; top, preliminary western blot of an expression tests shows double band pattern. Bottom, SDS-PAGE of a concentrated sample where glycosylations were stained using the periodic acid Schiff base (PAS) method B). Manually set up hanging drop crystallization results. Left: initial (1+1) µL droplet setup, at c = 2.0 mg mL^−1^. Right: refined (3+1) µL droplet setup, at c = 3.1 mg mL^−1^ C).

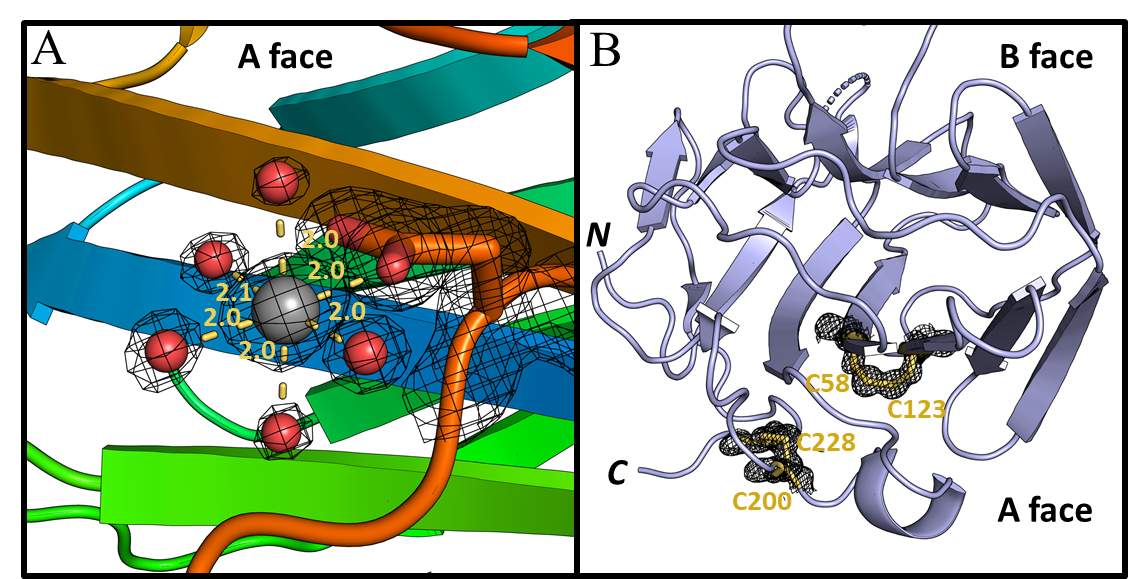

Fig. S2: Features of the ADGRG4-PTX structure. (2F_o_-F_c_)-type electron densities are presented at a contour level of 2.0 σ. Octahedral coordination sphere of Mg^2+^ (gray sphere) in front of the A face. Red spheres represent oxygen atoms of water molecules A). The magnesium ion was refined to an occupancy of 0.59 in the crystal structure. ADGRG4-PTX structure with the two disulfide bridges highlighted B).

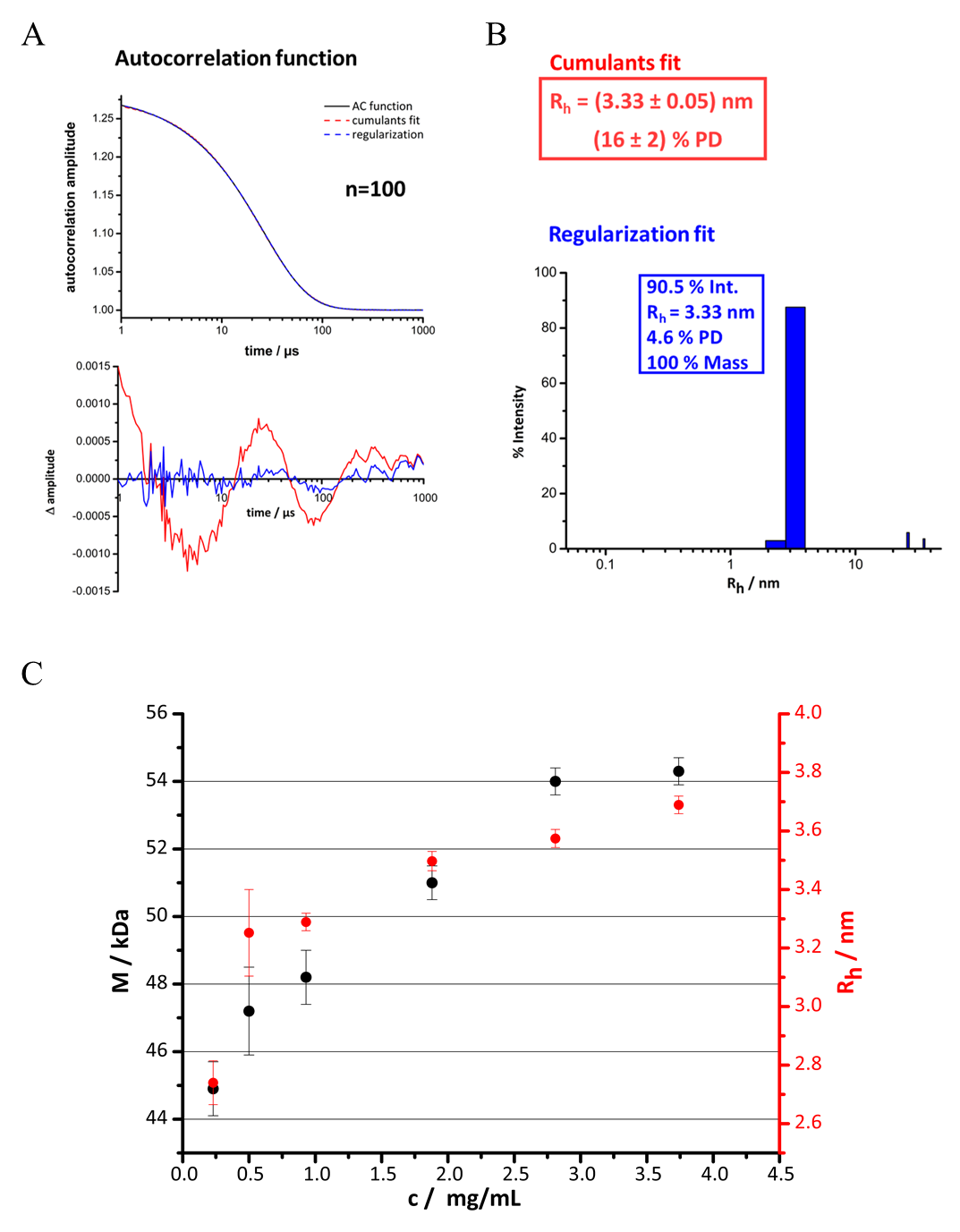

Fig. S3: Characterization of purified ADGRG4-PTX by light scattering. DLS of ADGRG4-PTX, measured at c = 1.0 mg mL^−1^ A). Results from the cumulants fit and regularization fit including the particle size distribution B). Dependence of the molecular mass (M, left ordinate, black points) as determined by SLS and the hydrodynamic radius (R_h_, right, red points) as determined by DLS on the protein concentration C). The measured samples were free of higher-order aggregates, as verified by DLS (compare S3A and B). All measurements (A-C) were performed at 20 °C. DLS was measured at an acquisition time of 5 s each. The displayed data are a mean result over 100 acquisitions for each concentration.

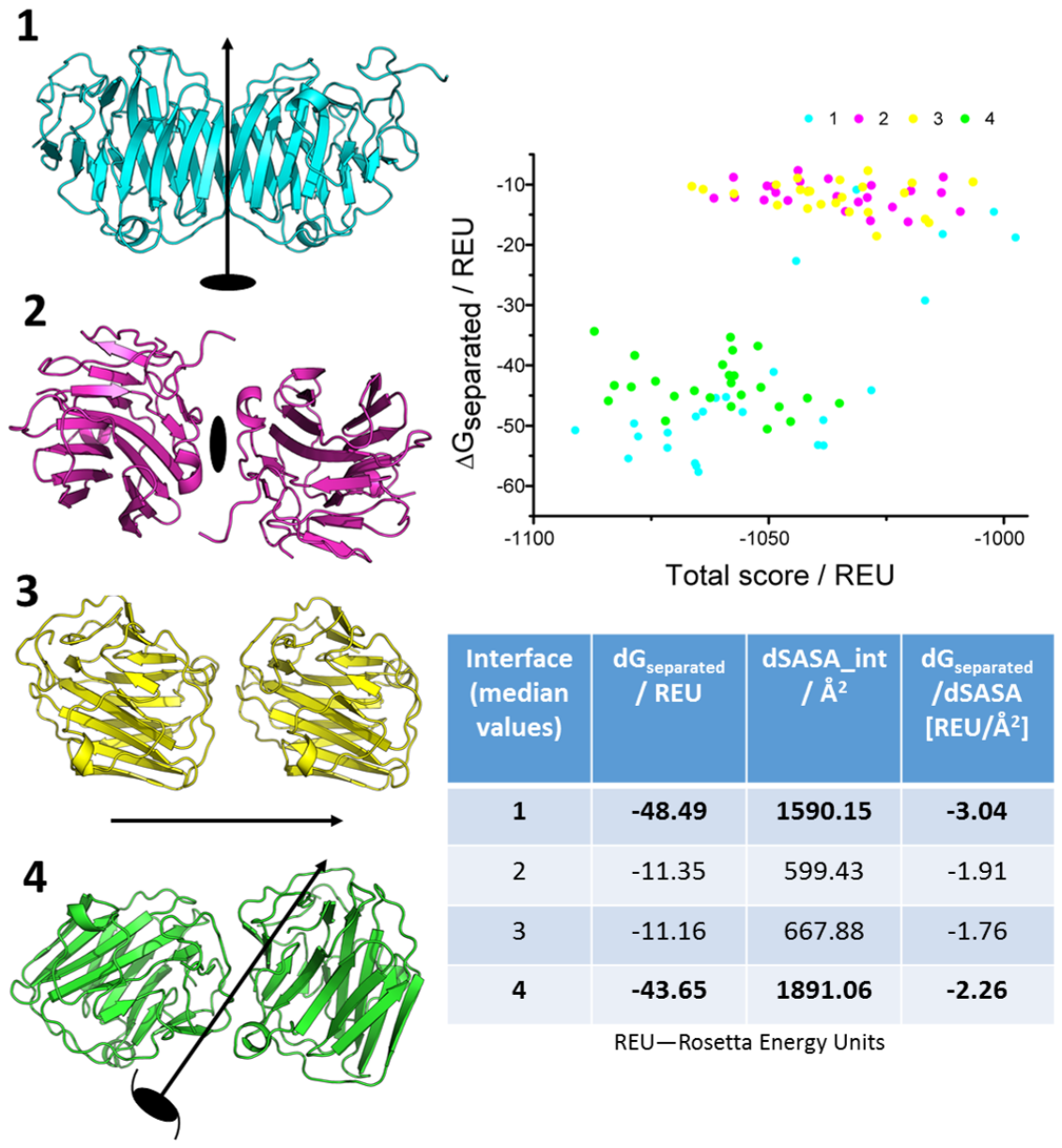

Fig. S4: Protein-protein interfaces present in the crystal lattice (*left*). The respective symmetry operators creating the symmetry mates are shown as black symbols. The right graph illustrates a ranking of the energetically relaxed homodimer structure models according to the total energy score in Rosetta energy units (REU, abscissa) as determined by Rosetta remodel and dG_separated_, which represents the free enthalpy difference due to dimer formation (ordinate). Each data point in the graph represents an individually relaxed dimer model. The table summarizes important interface measures by giving the respective median value of the 25 individual models each; dSASA_int: difference in solvent accessible surface area upon dimer separation (measure of dimer surface area) (Leaver-Fay et al. 2011).

A	B

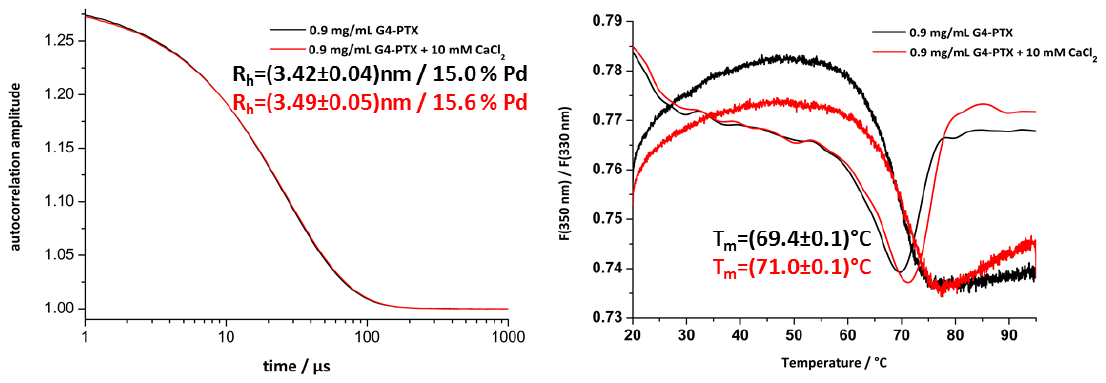

Fig. S5: Influence of Ca^2+^ on ADGRG4-PTX stability and hydrodynamic radius as determined by nanoDSF and DLS, respectively. DLS autocorrelation function shown for ADGRG4-PTX in SEC buffer and in presence of SEC buffer including10 mM CaCl_2_. The measurements were performed at 20 °C and an acquisition time of 5 s. The displayed data are a mean result over 100 acquisitions. R_h_-values and polydispersities are retrieved from the respective cumulant fits A). nanoDSF melting curves of ADGRG4-PTX in SEC buffer and in presence of additional 10 mM CaCl_2_. The data were averaged from three single measurements B).

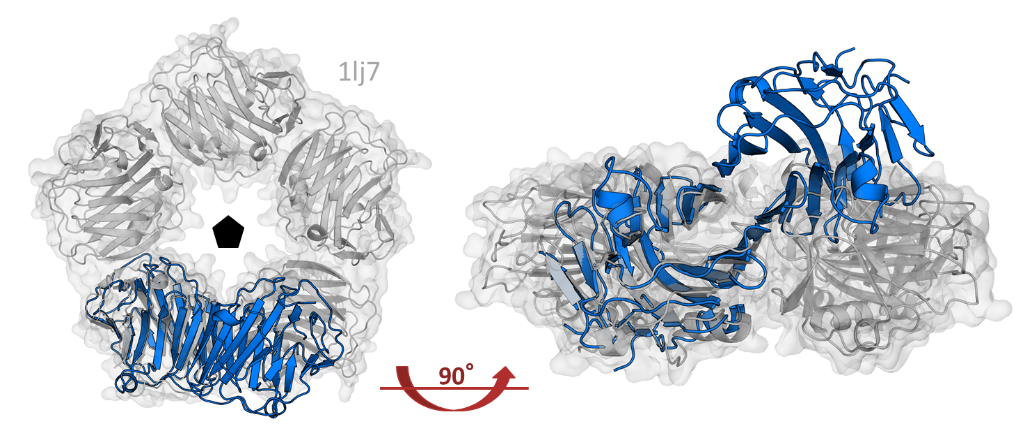

Fig. S6: Structural alignment of the ADGRG4-PTX dimer (blue) and the CRP pentamer (grey) based on the coordinates of one protomer.

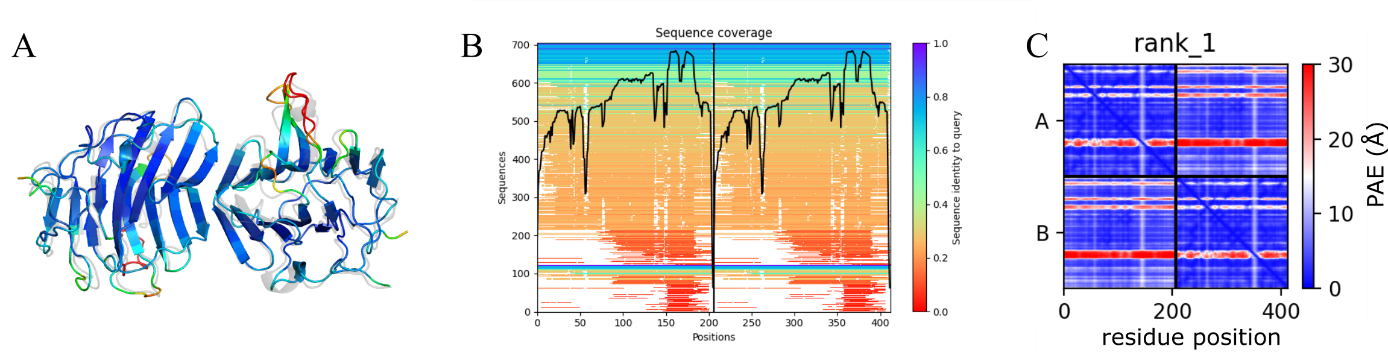

Fig. S7: Prediction of the ADGRG4-PTX dimer structure by AlphaFold (Jumper et al. 2021) as implemented in ColabFold (Mirdita et al. 2021). The predicted dimer structure is shown in rainbow colors according to the pLDDT value, a per-residue measure of local confidence on a scale from 0 - 100. Blue indicates high values (high confidence, maximum 100) and red low values (60 or lower). Shown in grey is the superimposed ADGRG4-PTX crystal structure. 78.2 % of the residues have high confidence (pLDDT > 90) and 14.6 % have medium confidence (50 < pLDDT ≤ 90) A). The sequence coverage specifies the number of sequences that contribute to modelling different regions of the protein B). The Predicted Aligned Error (PAE) reports the expected positional error at residue x, when the predicted and true structures are aligned on residue y. This value is useful to assess the confidence in the relative orientation of the protomers, i.e., in the dimer structure. In the upper left and lower right quadrants, we observe the positional errors of residues within the two domains of the dimer (average PAE = 5.4 Å). The upper right and lower left quadrants show the positional accuracy of the two protomers of the dimer relative to each other (average PAE = 8.8 Å). These low values indicate the high confidence in complex prediction, in agreement with the good match of the predicted and experimental dimer models (A) C).

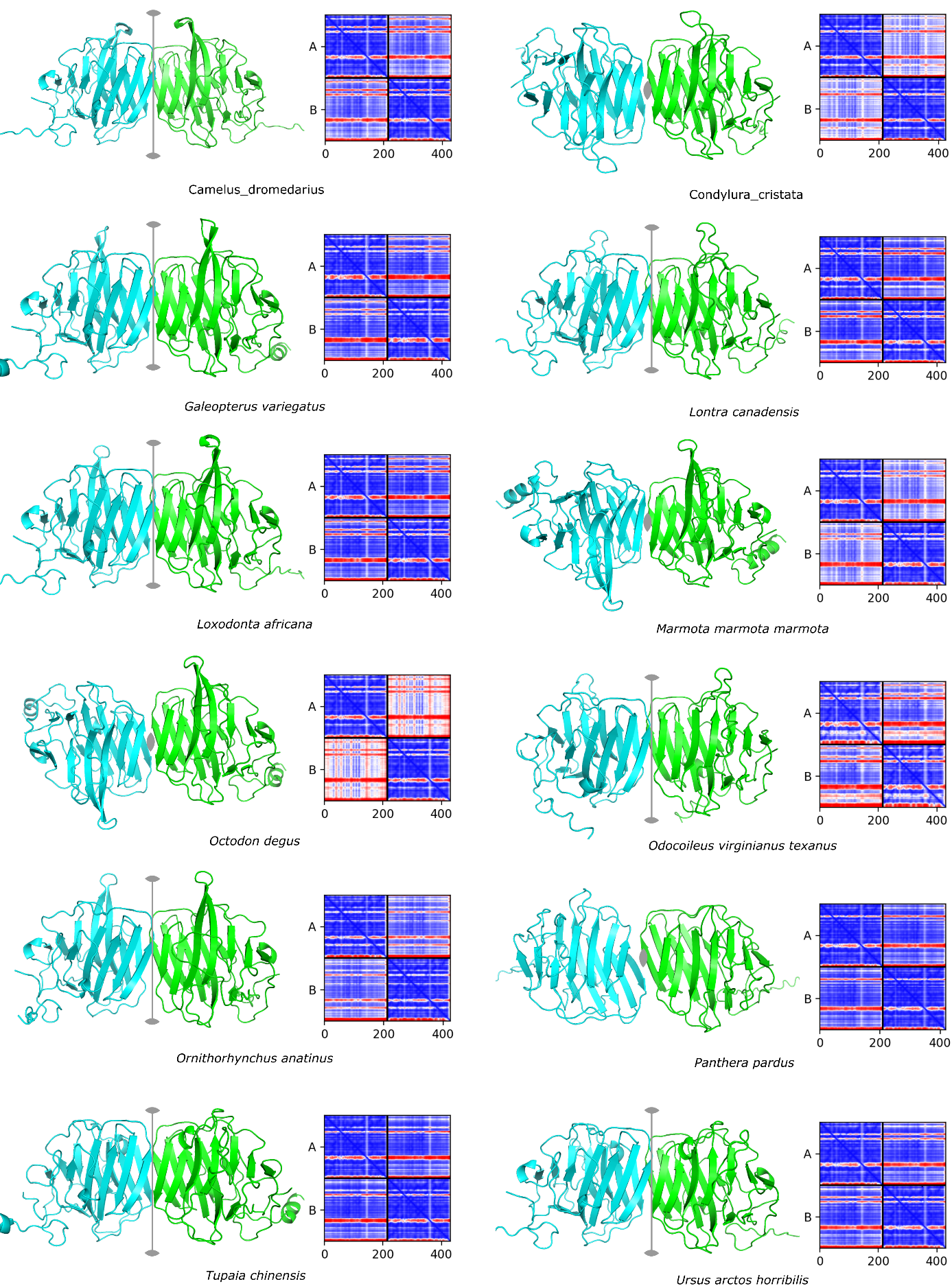

Figure S8: Prediction of the ADGRG4 pentraxin domain dimer structures for selected mammalian species by AlphaFold (Jumper et al. 2021) as implemented in ColabFold (Mirdita et al. 2021). Eight out of twelve dimers are predicted to form the same type of dimer as observed for the human ADGRG4 pentraxin domain. These have the molecular dimer axis running in the plane of the two sandwiched β-sheets. For the remaining four dimers, the same edge of the protomer is involved in dimer formation, but the corresponding β-strands of the two protomers (β7-β7' and β10-β10') directly interact to form the extended β-sheets. In this type of dimer, the two-fold axis is running perpendicular to the sandwiched β-sheets (see species *Condylura*, *Marmota*, *Octodon* and *Panthera*). For these predictions, the cross-diagonal quarters of the PAE plot (which indicate the confidence in dimer prediction) are lower compared to the other predictions. An exception is the prediction for the *Panthera pardus* G4 pentraxin domain. However, in this predicted structure two β-strands are too far apart for direct interaction. The two types of dimer probably generate similar co-evolution patterns for residues across the dimer interface as the same edge of the protomer is involved in dimer formation and the hydrogen-bonding interactions between the main-chain atoms of the interacting β-strands is mostly sequence-independent.

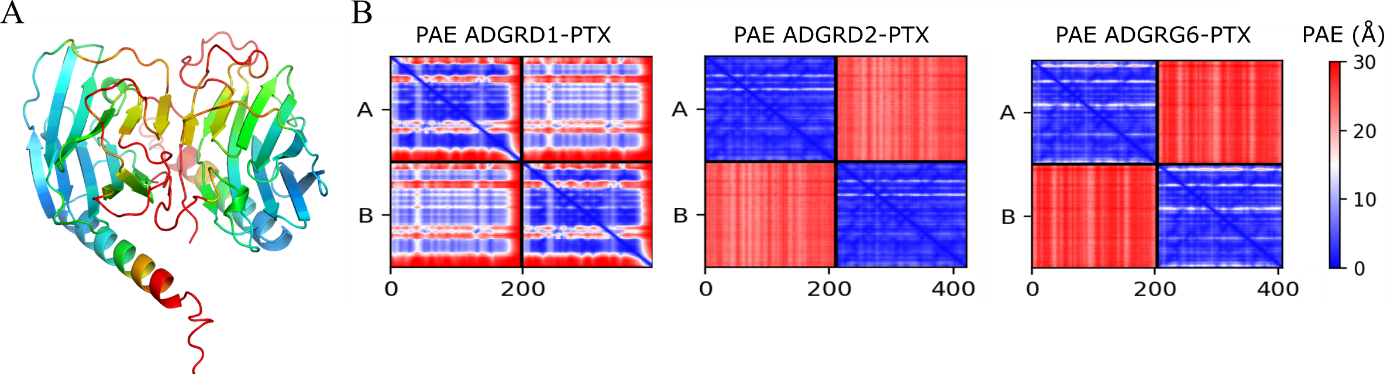

Fig. S9: AlphaFold dimer predictions of the pentraxin domains of ADGRD1, ADGRD2 and ADGRG6. Predicted dimer fold of ADGRD1. Colors indicate the pLDDT value as in Fig. S7A. The confidence (pLLDT) values for the protomer fold prediction are lower than for the other PTX-like domains of aGPCRs. The dimer interface is formed by the edges at the opposite site of the central β-sheet compared to ADGRG4. Furthermore, the regions involved in dimer formation have only medium or low confidence scores A). PAE values of the dimer predictions. The prediction indicates possible dimer formation for AGDRD1. For ADGRD2 and ADGRG6 no dimer structure could be predicted B).

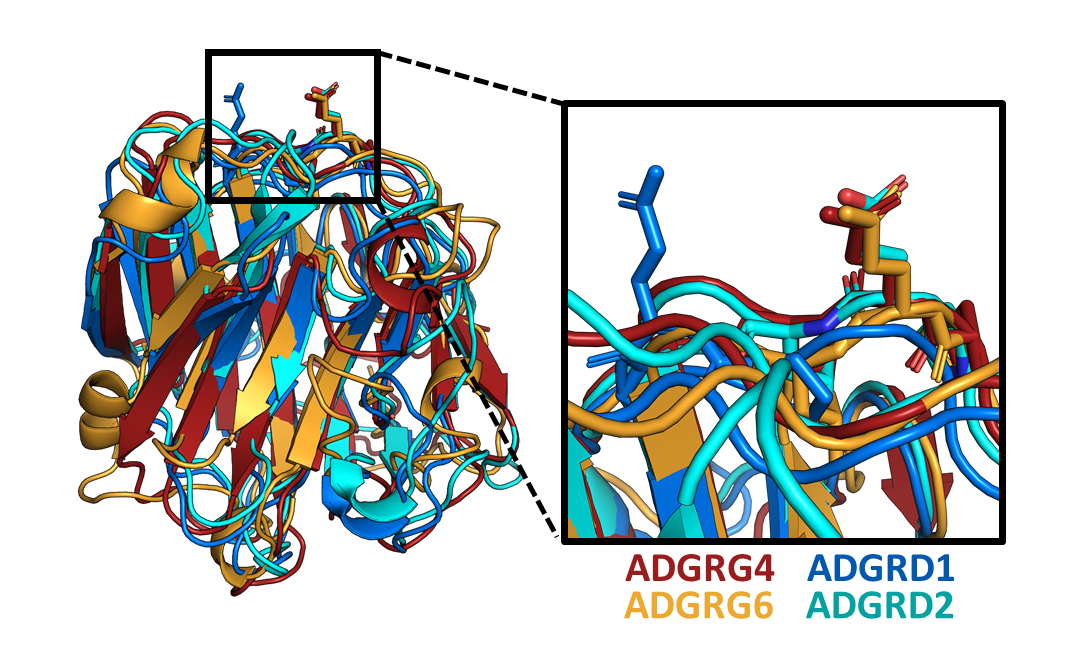

Fig. S10: Structural alignment of the four PTX-like domains of human ADGRD1, ADGRD2, ADGRG4, and ADGRG6, highlighting the conserved “PEL” motif. Except for ADGRG4-PTX, the other protein domain structures were predicted by homology modeling using the Phyre2 server in normal mode (Kelley et al. 2015). Note that the highly conserved glutamate residue protrudes from the surface and aligns nicely between the homologs, except for the PEG sequence in ADGRD1-PTX (compare Fig. 10).

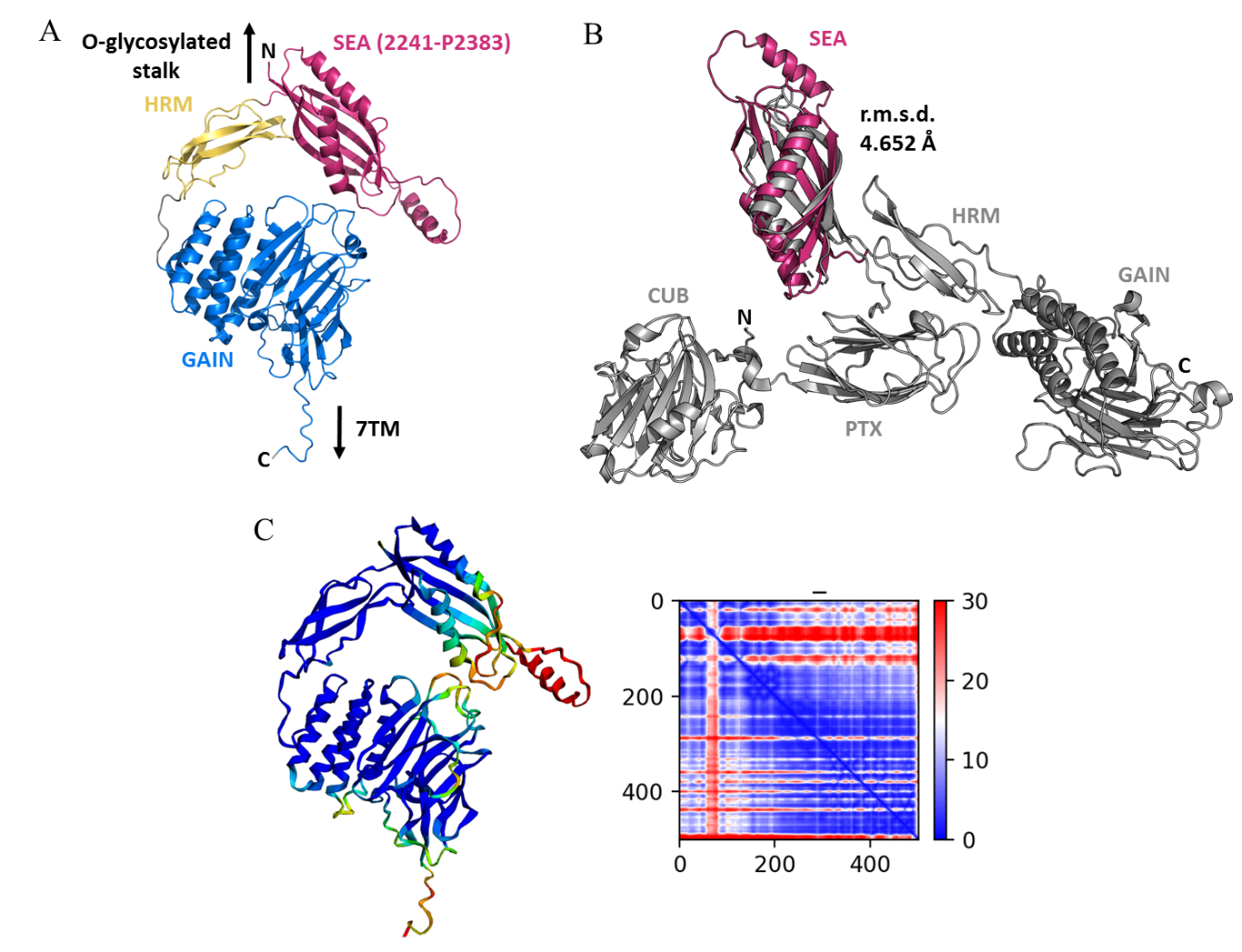

Fig. S11: AlphaFold prediction (Jumper et al. 2021; Leaver-Fay et al. 2011) of the non-mucin-like ADGRG4 ECR section close beside the 7TM. The three predicted protein domains are individually colored, the SEA domain spans from amino acids 2241–2383 and was not detectable by homology modeling A). Structural alignment of the SEA domains of the closely related ADGRG4 (*pink*) and ADGRG6 (*grey*, crystal structure: PDB 6v55) B). . pLDDT-values plotted onto the predicted ADGRG4 SEA-HRM-GAIN structure as in Fig. S7A and the respective PAE plot of the prediction C).

Tab. S1: Selection of closest structural relatives as found using PDBeFold (Krissinel und Henrick 2004).

| **PDB entry** | **Protein** | **r.m.s.d / Å** | **Seq. i.d. / %** |
| --- | --- | --- | --- |
| **6ype** | human neuronal  pentraxin (NP1)  PTX Domain | 1.66 | 25 |
| **1b09** | human CRP | 1.67 | 20 |
| **2a3y** | human SAP | 1.80 | 23 |
| **4pbo** | zebrafish short-chain pentraxin protein | 1.87 | 25 |
| **3flp** | *Limulus polyphemus* SAP-like pentraxin | 1.95 | 21 |
| **2rld** | Rat neurexin 1β | 2.12 | 12 |
| **3pve** | mouse G2 domain  of agrin | 2.35 | 14 |
| **3sh4** | Laminin G like domain from human perlecan | 2.40 | 10 |
| **2e6v** | *Canis lupus* VIP36 exoplasmic/lumenal domain | 2.41 | 7 |
| **4gkx** | human ERGIC-53 carbohydrate-binding domain | 2.42 | 10 |

Tab. S2: Crystallographic statistics of data collection and model refinement.

| **Data collection** | |
| --- | --- |
| Wavelength / Å | 0.9762 |
| Resolution range / Å | 51.71—1.36 (1.38—1.36) |
| Space group | C2 |
| a, b, c / Å | 113.12, 40.24, 54.91 |
| β / ° | 113.9 |
| Total reflections | 302484 (1468) |
| Unique reflections | 43935 (282) |
| Redundancy | 6.9 (5.2) |
| Completeness / % | 89.7 (19.6) |
| Mean I/σ(I) | 29.6 (2.9) |
| R_meas_ / % | 2.8 (52.8) |
| R_pim_ / % | 1.1 (22.5) |
| CC1/2 / % | 99.9 (91.7) |
| Wilson B-factor / Å^2^ | 19.0 |
| **Refinement statistics** | |
| Rwork / Rfree (%) | 15.4 (30.1) / 18.7 (42.3) |
| r.m.s.d. bonds (Å) / angles (°) | 0.01 / 1.063 |
| Clashscore | 3.43 |
| Ramachandran favored regions / % | 97.88 |
| Ramachandran outliers / % | 0.00 |
| Rotamer outliers / % | 0.56 |
| Number of non-hydrogen atoms | 1721 |
| Protein | 1613 |
| Heterogen | 1 |
| Solvent | 107 |

Tab. S3: Basic SAXS parameters of an ADGRG4-PTX concentration series. The R_g_ values were obtained from the Guinier approximation or the pair-distance distribution (values in parentheses) using PRIMUS. The p(r)-vs.-r-function also yielded D_max_, the longest dimension of the particle. Pair-distance distributions were optimized automatically in PRIMUS applying the autognom function.

| **c(G4-PTX) / mg mL^-1^** | **Guinier R_g_ (real-space) / Å** | **D_max_ / Å** |
| --- | --- | --- |
| 0.27 | 26.8 ± 1.5 (25.7) | 71.3 |
| 0.55 | 27.7 ± 1.9 (27.1) | 80.3 |
| 1.07 | 28.8 ± 3.2 (28.8) | 89.9 |
| 1.88 | 29.6 ± 4.9 (30.1) | 96.8 |
| 4.37 | 29.9 ± 5.7 (30.5) | 99.2 |
| 6.38 | 29.8 ± 2.9 (30.4) | 99.0 |
| 7.51 | 29.4 ± 2.0 (30.0) | 96.2 |
| 9.15 | 29.3 ± 0.1 (30.0) | 107.7 |
